# ProMaya: a hierarchical universal Deep Learning framework for accurate and interpretable Protein-Protein interaction identification

**DOI:** 10.64898/2026.04.03.716278

**Authors:** Umesh Bhati, Sagar Gupta, Veerbhan Kesarwani, Ravi Shankar

**Author notes:** Authors’ email addresses: UB.

## Abstract

Protein-protein interactions (PPIs) are molecular lego which define the physical states of cells. Accurately identifying PPIs remains challenging due to the interplay of several factors ranging from electrostatic to molecular geometry, topology, and physics. Existing computational approaches capture only fragments of this orchestra, limiting their generalizability across protein families and interaction types. Here, we present ProMaya, a hierarchical multi-scale Graph-transformer framework which has learned the 3D atomic geometry, electronic distribution, residue-level structure and disorder, surface mass-density signatures, and large protein language-model embeddings of interacting proteins. Highly comprehensively benchmarked across nine species and 47 GB experimentally validated data, ProMaya achieved consistently >95% average accuracy, outperforming state-of-the-art tools by >12%. Its in-built explainability provides clear mechanistic reasoning for the observed interactions. The first time introduced atomic and protein language learning has dramatically made it attain an outstanding level for PPI discovery in absolutely universal manner which can deal any species and potent to even bypass costly experiments.

## 1. Introduction

Proteins are fundamental macromolecules that play a pivotal role in the structure, function, and regulation of cell systems **[1]**. Composed of long chains of amino acids, proteins are synthesized based on genetic information encoded in DNA, and they perform a vast array of biological functions essential to life **[2]**. They operate within intricate cellular environments where their functions are mediated largely through physical associations with other molecules **[3]**. Rather than acting in isolation, the vast majority of proteins coordinate through complex interaction networks **[4]**, forming the physical existence of every biological process, from signal transduction to metabolic control, regulation, immune responses, and replication **[5-6]**. Mapping these protein-protein interactions (PPIs) are essential for understanding the ever dynamic cellular systems, enabling mechanistic insights into condition specific responses which will be enabling us to design suitable interventions **[5, 7]**. Yet despite their importance, current view of the interactome is far from complete. Even in model organisms such as *Homo sapiens, Saccharomyces cerevisiae*, and *Arabidopsis thaliana*, only a small proportion of all possible PPIs have been experimentally validated **[8-9]**. These gaps reflect both the biochemical complexity of protein association and the practical limitations of experimental techniques. The possible numbers of interactions in any biological system is not pragmatic enough to be captured by experiments, even for a single state and moment.

Classical experimental approaches including yeast two-hybrid (Y2H) **[10]**, tandem affinity purification-mass spectrometry (TAP-MS) **[11]**, co-immunoprecipitation, protein microarrays, and low-throughput biophysical assays, remain indispensable for interactome identification but are inherently constrained **(Figure 1B) [12-13]**. They often require large protein quantities, rely heavily on prior knowledge such as available antibodies, and suffer from high false-positive rates, especially when examining transient, weak, or context-specific interactions. These techniques are often expensive, technically demanding, and labor-intensive to implement. Large-scale interactome mapping experiments typically require extensive cloning, optimized expression systems, high-quality antibodies, and careful experimental design. As a result, despite decades of experimental effort, generating comprehensive interactome maps remains slow and resource-intensive process, missing majority of interaction space.

**Figure 1:**
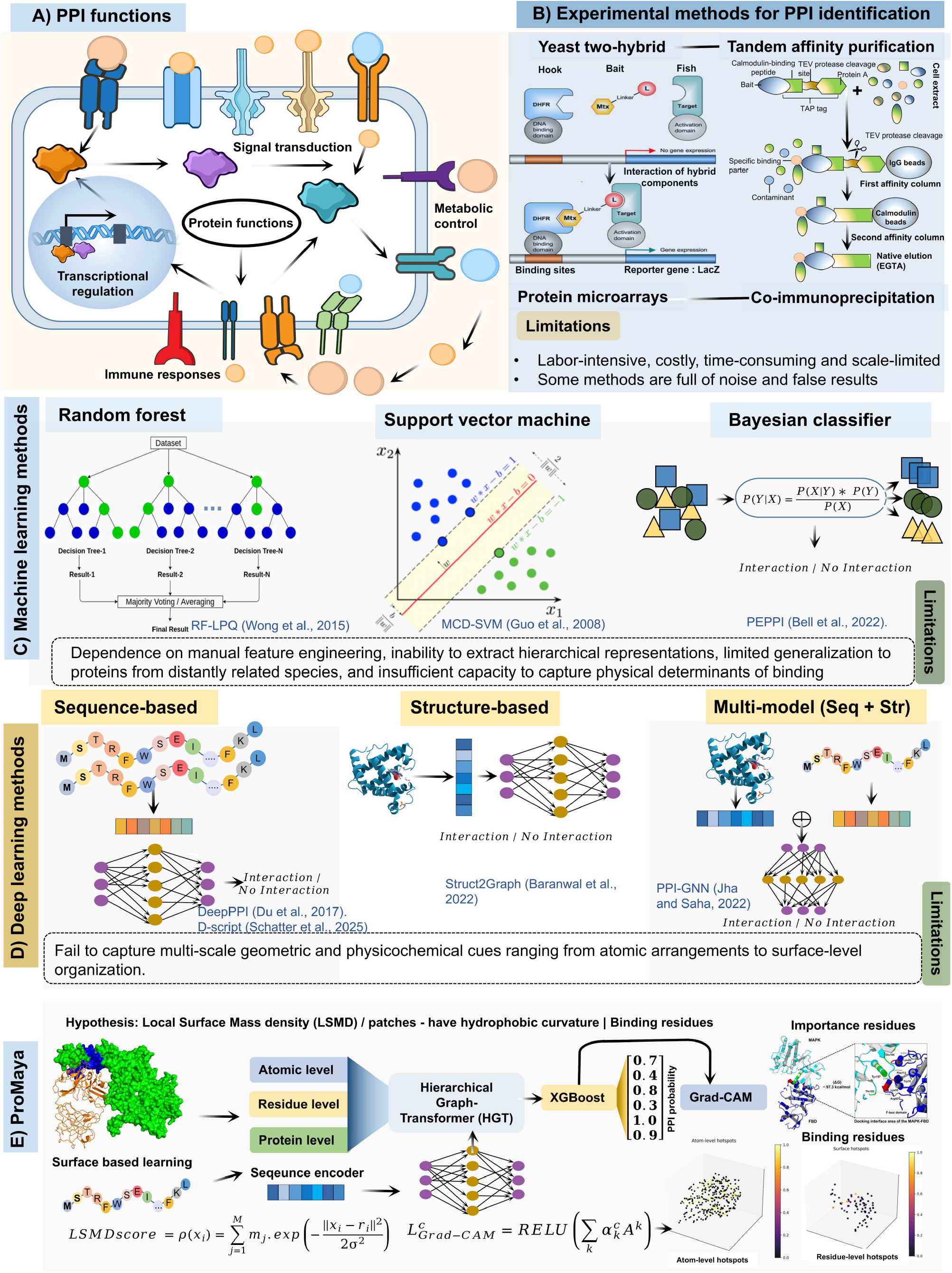
Overview of protein–protein interaction (PPI) biology. **(A)** Biological roles of protein– protein interactions in cellular systems. PPIs regulate essential cellular processes including signal transduction, transcriptional regulation, metabolic control, and immune responses, highlighting their central role in maintaining cellular homeostasis. **(B)** Experimental approaches for PPI identification. Representative methods include yeast two-hybrid (Y2H), tandem affinity purification coupled with mass spectrometry (TAP–MS), protein microarrays, and co-immunoprecipitation. Although widely used, these approaches are labor-intensive, costly, time-consuming, and often susceptible to experimental noise and false positives, limiting large-scale interactome mapping. **(C)** Classical machine-learning approaches for computational PPI prediction. Methods such as random forests, support vector machines (SVM), and Bayesian classifiers utilize handcrafted sequence-derived features to classify interacting and non-interacting protein pairs. However, these models rely heavily on manual feature engineering and exhibit limited generalization across diverse protein families. **(D)** Deep-learning approaches for PPI prediction. These approaches can be broadly categorized into sequence-based models, structure-based models, and multimodal frameworks integrating sequence and structural information. Despite improved predictive performance, these methods often fail to capture multi-scale geometric and physicochemical determinants of binding interfaces. **(E)** Overview of the ProMaya framework. ProMaya models protein interactions through hierarchical multimodal representations integrating atomic-, residue-, and protein-level features derived from surface-based geometric learning and local surface mass density (LSMD). These features are processed using a hierarchical graph transformer (HGT). Grad-CAM-based interpretability highlights key interface residues contributing to predicted PPIs, enabling mechanistic insights into binding determinants.

At the same time, advances in next-generation sequencing (NGS) have revolutionized molecular biology by dramatically increasing the volume of genomic and transcriptomic data available for nearly all organisms **[14-15]**. The rapid decline in sequencing costs and improvements in high-throughput technologies have enabled the generation of massive genomics datasets. While these developments have produced unprecedented insight into genomics, a comparable expansion of protein-centric datasets, such as proteomic measurements and interaction networks, has been far more limited **[16]**. Unlike nucleic acids, proteins exhibit complex structural dynamics, post-translational modifications, and context-dependent interactions that are difficult to measure experimentally at scale. Consequently, although sequence data for millions of proteins are now available, the experimental characterization of their structures and interaction partners has not kept the pace **[17]**. This imbalance between rapidly accumulating sequence data and comparatively scarce structural and interactome information has produced what is commonly described as the sequence-structure-interaction gap.

These challenges have motivated the development of computational approaches for PPI discovery **[18]**. Early computational PPI models predominantly relied on classical machine learning (ML) techniques applied to handcrafted protein features. Amino acid composition, physicochemical properties, evolutionary substitution scores, domain co-occurrence, and secondary structure propensities based features were commonly engineered to construct feature vectors for ML classifiers. Among the most influential methods were support vector machines (SVM), random forest (RF), and Bayesian classifiers such as PEPPI **[19]**. These models demonstrated that sequence-derived descriptors contain sufficient information to discover certain classes of interactions, and they provided foundational insights into computational PPI inference. However, they also suffered from serious limitations: dependence on manual feature engineering, inability to extract hierarchical graph representations, limited generalization to proteins from distantly related species, and insufficient capacity to capture physical determinants of binding **[20]**. Their biggest limitation was that they were not able to capture more informative intermediate variables, something which only deep learning (DL) could achieve.

Subsequently, DL has revolutionized computational biology by enabling automatic, data-driven hidden and meta features derivations from raw sequences, structures, and molecular graphs **[21]**. DL-based PPI algorithms can be seen forming three major categories: based on sequence, structure, and multimodal fusion **[22]**:

i. **Sequence-based approaches** remain the most widespread due to the abundance/availability of protein sequence data and the computational scalability required for proteome-wide analyses like classification of proteins. Models such as DeepPPI **[23]** and D-script **[24]** leverage recurrent neural networks (RNN), convolutional architectures, and transformer-based encoders to identify sequence patterns necessary to identify PPIs. Very recently, embeddings from protein large language models (LLMs) **[25-27]** have significantly improved representational richness by capturing evolutionary constraints, mutational co-variation, and functional signatures directly from protein sequence corpora. However, sequence-based models inherently lack explicit structural information such as geometric complementarity, surface topology, and spatial arrangement of interaction residues which are fundamental determinants of physical binding. As a result, these methods often struggle to distinguish proteins that share similar sequence profiles but differ in structural compatibility.
ii. **Structure-based approaches** address this limitation by incorporating three-dimensional (3D) information. The advent of AlphaFold2 **[28]** has dramatically expanded structural coverage, enabling the development of graph-based models such as PPI-GNN **[29]** and Struct2Graph **[30]**, which leverage geometric and topological features to identify interaction interfaces. These methods improve the modeling of residue proximity and surface characteristics; however, they remain constrained by incomplete structural coverage, conformational variability, and an inability to capture transient or flexible interactions. Notably, intrinsically disordered regions (IDRs), which play a central role in many PPIs, are poorly represented in conventional structure-centric frameworks, limiting their applicability across diverse interaction types **[31]**.
iii. **Multimodal fusion based approaches** have emerged to integrate heterogeneous data sources covering sequence and structural properties together. These models employ sequence and structure information together to synergize evolutionary, structural, physicochemical, and geometric information into unified representations more consistent with biological principles. For example, Jha et. al., (2022) **[29]** developed a multimodal PPI framework employing 3D protein volumes processed by ResNet50 alongside conjoint triad and autocovariance sequence representations **[29]**. Similarly, recent studies have incorporated distance maps, energy-based descriptors, and topology-aware graph embeddings **[32-37]**. While these multimodal approaches outperform unimodal variants, they still fall short in capturing the nuanced, partner-specific interaction patterns required for high-resolution PPI identification. Many GNN-based architectures primarily aggregate neighborhood features rather than explicitly modeling the relational dependencies unique to each protein pair, limiting both its accuracy and generalization capability across proteome spanning multiple species **(Figure 1C)**.

Despite all these development, three critical limitations persist across existing methodologies. First, unimodal architectures (sequence or structure) inadequately capture hierarchical dependencies, Residue interactions are modulated not only by local contacts but also by global protein topology and dynamics. Second, no framework explicitly models local surface mass density (LSMD), the spatial distribution of atomic mass across protein surfaces, as a physical determinant of binding. LSMD governs van der Waals forces and hydrophobic collapse, which yet remains unquantified in PPI prediction. Third, geometric representations lack multi-scale resolution, failing to reconcile atomic-level physio-chemical details with mesoscale interface morphology **[38]**. Consequently, even state-of-the-art (SOTA) tools such as PPI-GNN, exhibit lower accuracy for transient interactions and less application when it comes to cross-species generalizability.

Motivated by all these major bottlenecks, here we introduce ProMaya, a deep learning framework that redefines multimodal PPI discovery by explicitly modeling LSMD as a primary driver of molecular recognition while simultaneously capturing the dynamic contributions of IDRs to protein binding. Our core hypothesis is that protein–protein interaction is governed by complementary “mass-density fingerprints” on opposing surfaces dense regions arising from tightly packed hydrophobic cores, π-stacking aromatic clusters, and buried polar networks create distinctive physicochemical signatures that guide partner selection more effectively than electrostatics or hydrophobicity alone. However, a substantial subset of interactions particularly in signaling and regulatory networks are mediated not by rigid interfaces but by flexible, disorder-enriched segments that undergo disorder-to-order transitions, engaging via short linear motifs **[29, 39]**. These IDRs exhibit low LSMD, high conformational entropy, and unique sequence signatures that cannot be captured by geometry or structure alone. To account for these dual modes of molecular recognition, ProMaya integrates mass-weighted surface point clouds, atomic-resolution geometric graphs, and IDR propensity scores, unifying them with evolutionary constraints derived from large protein language models such as ProtTrans **[27]**. This combination enables ProMaya to accurately model both structured binding interfaces and flexible disorder-mediated interactions, providing a holistic, multi-scale representation of the determinants that govern PPI specificity across proteome, in a completely universal manner.

To implement this principle, ProMaya synergizes three innovations:

**1) Multi-scale geometric learning:** atomic coordinates were transformed into a Pointnet++ derived **[38]** point cloud where each vertex encodes LSMD alongside electrostatic, curvature, and chemical features. **2) Hierarchical graph transformers (HGT):** a novel HGT architecture processes features across atomic (Å-scale), residue (nm-scale), and protein (µm-scale) resolutions, with cross-level attention mechanisms that propagate physical constraints upward and functional context downward. **3) Multimodal sequence-structure fusion:** evolutionary signals from ProtTrans embeddings **[27]** were integrated with geometric features via gated cross-attention, preserving biological context, otherwise lost in pure structure-based models. **Figure 3** provides full implementation details of ProMaya architecture.

ProMaya achieved remarkable accuracy and consistency while surpassing 97% accuracy, with average accuracy of >95% when tested across a huge volume of experimental data covering different species. It outperformed SOTA methods such as PPI-GNN **[29]** (80.9%), D-SCRIPT **[24]** (83.4%), DeepPPI **[23]** (81.3%), and PEPPI **[19]** (78.8%) with big margin when benchmarked across four independent datasets. ProMaya achieves, to our knowledge, the strongest cross-species and cold-start performance reported for PPI identification, across nine species covering major taxonomical groups and with 12% gains over the existing state-of-the-art methods. This also make it a strong alternative to costly and cumbersome experiments, while holding the promise to accelerate the field of systems biology.

## 2. Materials and methods

### 2.1 Dataset preparation

Experimentally determined binary protein–protein complexes were sourced from PDBbind 2021 **[40]** and DIPS-PLUS **[41]**, comprising an initial pool of 46,706 structures across six model organisms (*Homo sapiens, Saccharomyces cerevisiae, Escherichia coli, Arabidopsis thaliana, Drosophila melanogaster*, and *Caenorhabditis elegans*) **(Supplementary Table S1 Sheet 1-2)**. Structural quality was enforced by excluding entries with crystallographic resolution >3.0 Å, incomplete backbone chains, >5 consecutive missing residues, or non-physiological oligomeric states. Interface quality was quantified using the buried solvent-accessible surface area (SASA; ΔA_int_), calculated via FreeSASA **[42]** with a probe radius of 1.4 Å. Complexes were retained only if ΔA_int_ ≥ 200 Å^2^ and each chain contributed ≥7 interface residues (defined as ΔSASA>1 Å^2^ and ≥1 heavy atom within 5 Å of the partner) **(Figure 2B-C)**. Redundancy was removed by clustering all protein chains at 40% sequence identity using CD-HIT **[43]** (word length 2, 0.6, global identity mode). At most one representative complex per cluster-pair combination was retained, ensuring all chains within a CD-HIT cluster were confined exclusively to a single data split to prevent sequence-level leakage **(Supplementary Figure S1)**.

**Figure 2:**
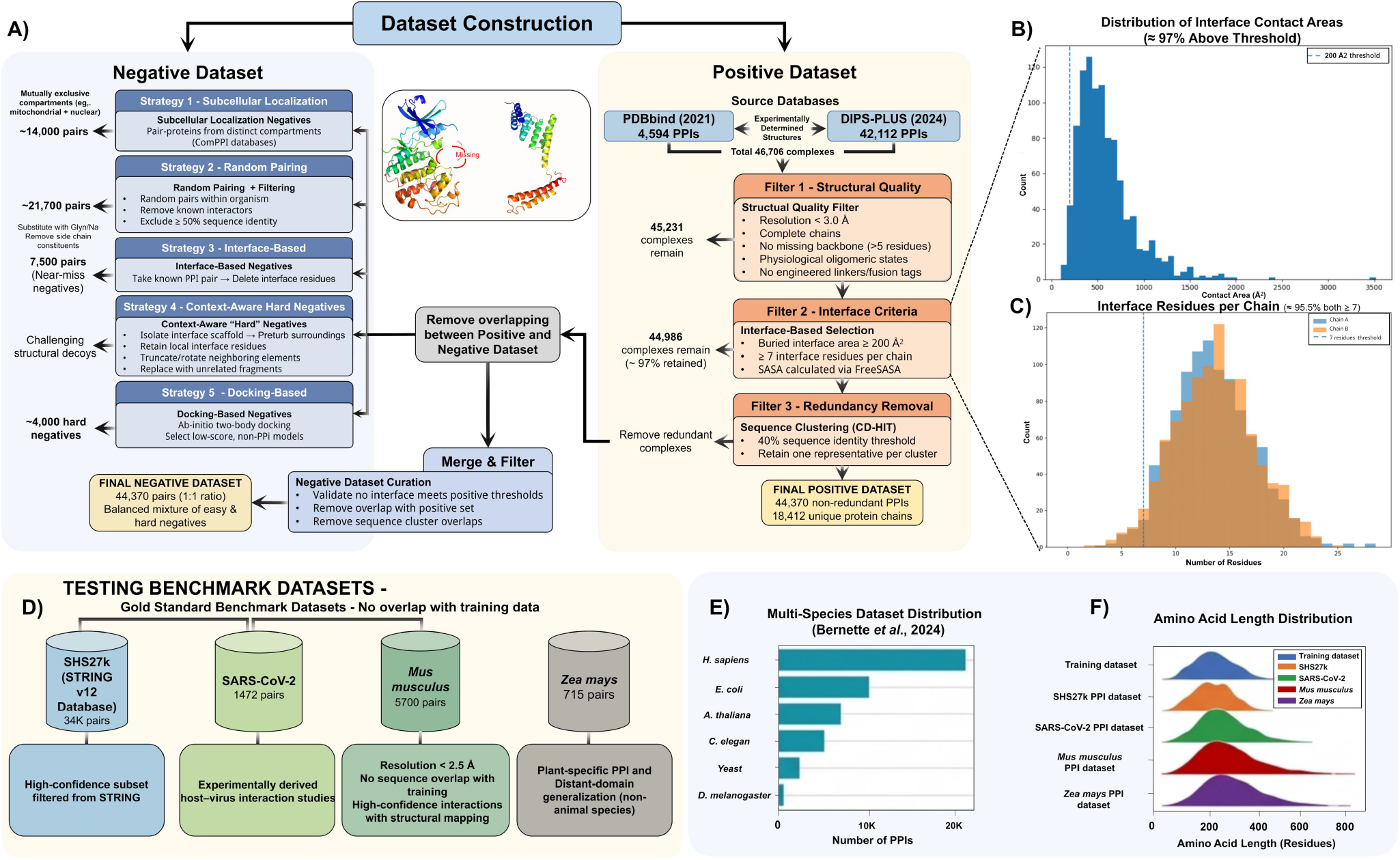
Construction and validation of the ProMaya protein–protein interaction dataset. **(A)** Dataset construction workflow. Positive PPI complexes were collected from experimentally validated databases (PDBbind and DIPS-PLUS) and Negative samples were generated using multiple complementary strategies. **(B)** Distribution of interface contact areas for positive complexes after filtering. The majority of complexes exceed the interface threshold (200 Å ^2^), ensuring biologically meaningful interaction surfaces. **(C)** Distribution of interface residues per chain. Most complexes contain more than seven interface residues per chain, confirming that the dataset predominantly contains well-defined interaction interfaces. **(D)** Independent benchmark datasets used for external validation. Gold-standard datasets including SHS27k, SARS-CoV-2, *Mus musculus*, and *Zea mays* were selected to ensure cross-species and cross-family evaluation without overlap with the training dataset. **(E)** Multi-species distribution of PPIs in the benchmark datasets, illustrating the diversity of organisms represented. **(F)** Amino acid length distributions across the training dataset and benchmark datasets, demonstrating comparable protein length characteristics and ensuring that the evaluation datasets represent realistic structural diversity.

**Figure 3:**
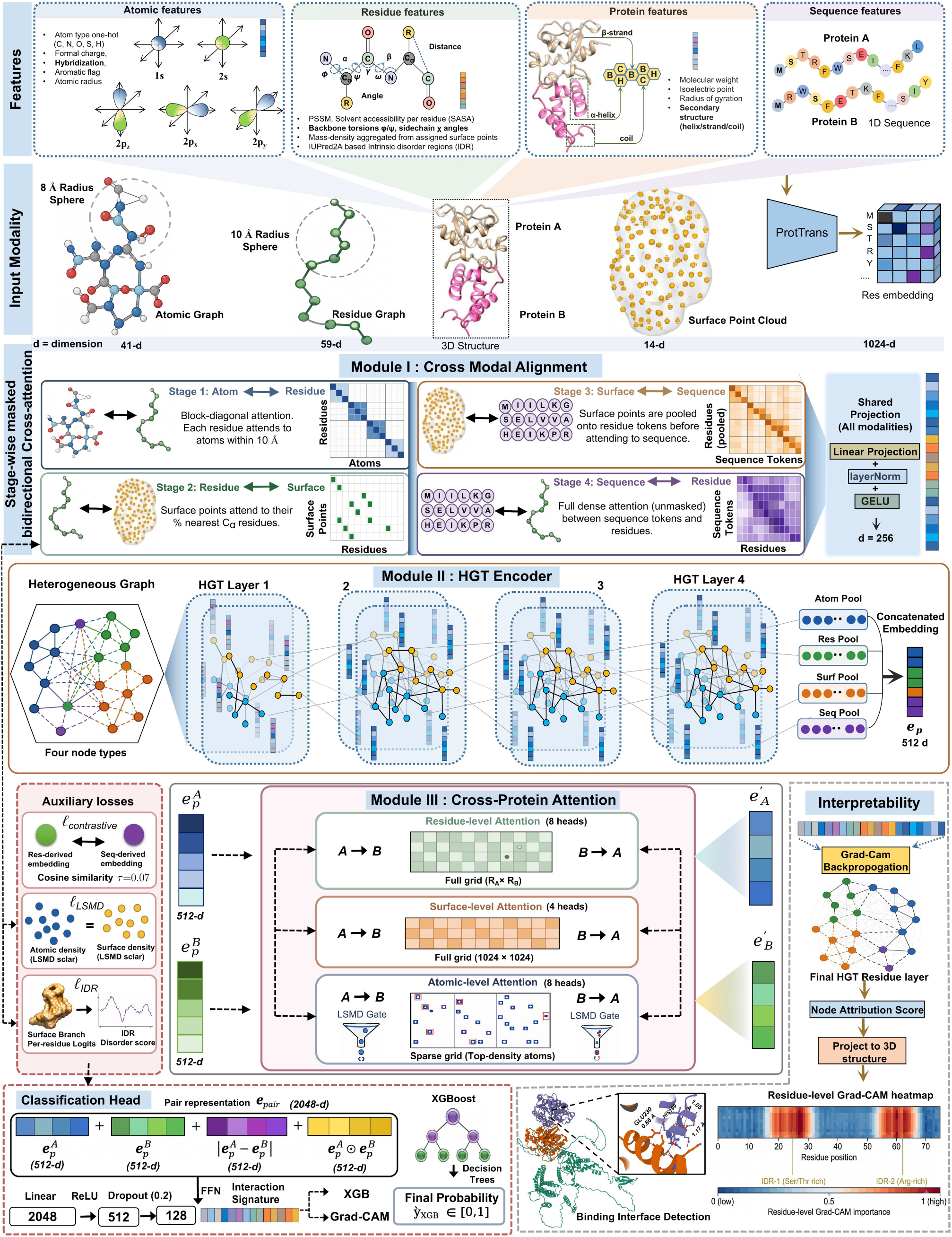
Architecture of ProMaya for multimodal protein–protein interaction identification. ProMaya integrates atomic, residue, protein, and sequence features through three major modules: cross-modal alignment, heterogeneous graph transformer (HGT) encoding, and cross-protein attention. Stage-wise masked cross-attention aligns representations across modalities, followed by HGT-based hierarchical message passing and pooling. Cross-protein attention captures residue-, surface-, and atom-level interaction patterns. The fused representations are classified using an XGBoost-based prediction head. Grad-CAM provides interpretable residue-level attribution and maps predicted interaction regions onto the 3D protein structures.

#### 2.1.1 Negative dataset formation

A composite negative dataset was constructed to match the positive set size (1:1 ratio) using five biologically motivated categories: **(i)** Subcellular-localization negatives (∼14,000 pairs), pairing proteins from mutually exclusive compartments (ComPPI **[44]**, confidence ≥0.5); **(ii)** Random-pairing negatives (∼21,700 pairs), intra-organism couplings excluding known interactors and pairs with sequence identity >50%; **(iii)** Interface-ablated negatives (∼7,500 pairs), generated by replacing interface residues with glycine/alanine and deleting side-chain coordinates; **(iv)** Context-aware hard negatives, retaining interface scaffolds while grafting structurally dissimilar fragments (SCOPe **[45]**, TM-score <0.4) into flanking regions; and **(v)** Docking-derived negatives (∼4,000 pairs), selected as low-scoring HADDOCK **[46]** decoys from *ab initio* docking of non-interacting pairs. All negatives were validated to violate positive-set interface thresholds and exhibit no overlap with positive chains or interface geometries **(Full detailed in Supplementary File Methods; Supplementary Table S1 Sheet 3)**.

#### 2.1.2 External benchmark datasets

Four independent external datasets, none overlapping with the training corpus, were used exclusively for generalization evaluation.

i. **SHS27k (STRING v12) :** Approximately 34,000 high-confidence human PPI pairs were retrieved from the STRING v12 database **[47]** with combined scores ≥ 700, filtered to retain only experimentally validated evidence channels. This benchmark enables direct comparison with published sequence-based and structure-based PPI tools **(Supplementary Table S1 Sheet 4)**.
ii. **SARS-CoV-2 host-pathogen dataset:** 1,472 experimentally determined SARS-CoV-2 viral protein–human host factor interaction pairs derived from AP-MS10 and cryo-EM structural studies **[48]**. All viral proteins in this set were absent from the training corpus.
iii. ***Mus musculus* PPI dataset:** 5,700 mouse PPI pairs resolved at crystallographic resolution < 2.5 Å, with no training-set sequence overlap at the 40% identity threshold **[49]**. High-confidence interactions were mapped to structural coordinates via PDBe-KB.
iv. ***Zea mays* PPI dataset:** 715 maize PPI pairs representing plant-specific interaction architectures including photosynthetic complexes and hormone signaling assemblies **[50]**. Sequence identity to training-set chains does not exceed 30% in >92% of cases **(Supplementary Table S1 Sheet 5)**.

### 2.2 Preliminaries and Problem Statement

We treat PPI prediction as a pairwise binary classification task over a structural protein population, distinct from the transductive PPI-network setting used by graph-propagation methods. Given a set of proteins *P* ={*P*_1_,…,*P*_N_}, each available as an experimentally resolved or computationally predicted 3-D structure with sequence *S*_*i*_, and a labeled training set *D*_*train*_ = {(*P*_*i*_ , *P*_*j*_, *y*_*ij*_)} with *y*_*ij*_ ∈ {0,1} , the objective is to learn a function

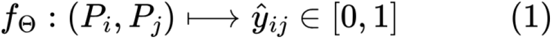

that generalizes to unseen pairs (*P*_*k*_, *P*_*l*_) ∉ *D*_*train*_, including pairs where neither *P*_*k*_ nor *P*_*l*_ was observed in any training pair (the Neither-Seen / cross-species regime). ProMaya consumes no PPI-network adjacency structure at inference time; consequently, it makes no assumption that the query pair is embedded in a previously seen interaction graph.

#### Input representation

Each protein *P* is represented simultaneously in four modalities:

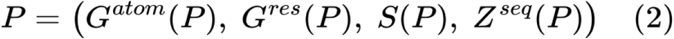

where *G*^*atom*^ (*P*) *G*^*res*^(*P*) and are the atomic and residue graphs **(Section 2.3)**, *S*(*P*) is the surface point cloud, and *Z*^*seq*^(*P*) ∈ ℝ^*L*×1024^ is the frozen ProtT5 embedding. ProMaya requires all four modalities to be populated. Crucially and this is the central departure from concatenation-style multimodal models it explicitly constrains them to be mutually consistent before fusion **(Section 2.4)**.

### 2.3 Multi-Scale Feature Extraction

#### 2.3.1 Atomic Graph and Local Surface Mass Density

The atomic graph is *G*^*atom*^ = (*V*^*atom*^, *E*^*atom*^) with

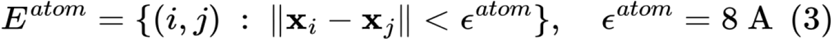

Hydrogens are excluded from *V*^*atom*^. Each node carries 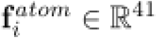, formed by concatenating an element one-hot, heavy-atom-type one-hot, Gasteiger partial charge, hybridization one-hot, van der Waals radius, backbone aromatic H-bond-donor-acceptor flags, chirality one-hot, a surface-exposure flag, and the LSMD scalar defined next.

##### LSMD

ProMaya’s central structural hypothesis is that a single physically transparent scalar, Gaussian-smoothed local atomic mass density, summarizes the net effect of several weak non-covalent forces that jointly stabilize an interface:

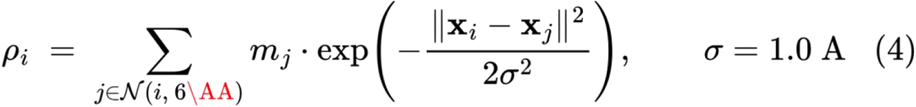

where *m*_*j*_ is the standard atomic weight of neighbor *j* and *N*(*i*,6Å) is the set of heavy atoms within 6 Å of atom . Because *σ* = 1.0 Å, the Gaussian kernel decays to <10^−8^ at the 6 Å boundary; the cutoff therefore serves as a hard upper bound for computational efficiency, while the effective support of the kernel is concentrated within ∼3 Å. *ρ*_*i*_ is z-scored per protein (removing size confounds) before use. Residue-level LSMD is obtained by mean aggregation,

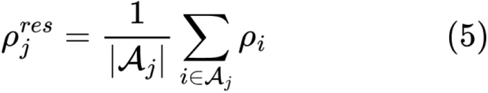

where *A*_*j*_ is the set of heavy atoms belonging to residue *j*. Surface-level LSMD is obtained by inverse-distance interpolation from the five nearest heavy atoms,

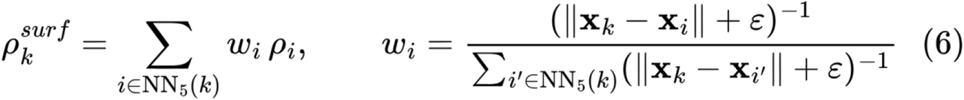

where NN_5_(*k*) denotes the five nearest heavy atoms to surface point *k* and *ε* = 10^−6^ Å prevents division by zero.

Atomic edges carry a 35-dimensional feature: 16-bin RBF encoding of ‖x_*i*_ − x_*j*_‖ over 0–8 Å; 8-bin polar and 8-bin azimuthal angles relative to the local *N*−*C*_*α*_−*C* backbone frame; and a 3-dimensional torsion-like dihedral between bonded heavy-atom planes.

#### 2.3.2 Residue Graph

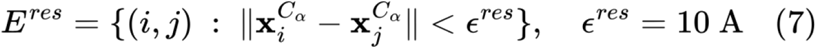

Each residue node carries 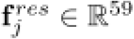:

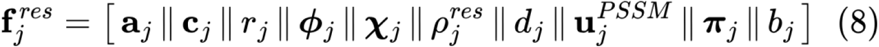

where **a**_*j*_ ∈ {0, 1}^21^ is the amino-acid one-hot; **c**_*j*_ ∈ {0, 1}^3^ the DSSP 3-state secondary-structure one-hot; *r*_*j*_ ∈[0, 1] the relative SASA; *ϕ*_*j*_ = (sin*ϕ*, cos*ϕ*, sin*ψ*, cos*ψ*) the backbone torsion encoding; ***χ***_*j*_ = (sin_*χ*1_, cos_*χ1*_) the side-chain torsion encoding; 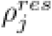 from Eq. (5); *d*_*j*_ ∈ [0, 1] the IUPred2A long-mode disorder score; 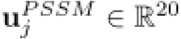 the PSI-BLAST position-specific scoring vector; π ∈ ℝ^5^ the physicochemical descriptor (hydropathy, polarity, volume, charge, molecular weight); and *b*_*j*_ ∈ {0, 1} the no-MSA indicator, set to 1 when PSI-BLAST returns fewer than three hits at *E* < 0.001, in which case 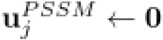.

Residue edges carry a 36-dimensional feature: 16-bin RBF of *C*_*α*_ −*C*_*α*_ distance; log-transformed sequence separation; backbone-hydrogen-bond indicator; same-secondary-structure-element indicator; 4 sequence-separation-range bins; and 10 relative-orientation descriptors.

#### 2.3.3 Surface Point Cloud

The solvent-accessible surface is computed by MSMS (probe radius 1.4 Å, density 3.0 Å ^−2^). The raw MSMS vertex set is downsampled to a fixed cardinality of *M* =1,024 points via iterative farthest-point sampling, ensuring that all proteins feed the surface encoder at identical tensor shapes regardless of physical size. Each retained point 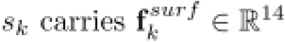 : Cartesian coordinate, surface normal, mean and Gaussian curvature, Poisson–Boltzmann electrostatic potential (APBS/PDB2PQR, pH 7.0), interpolated relative SASA, 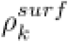 from Eq. (6), shape index, curvedness, and distance to the nearest backbone atom. A three-layer PointNet++ encoder with ball-query radii {5, 6, 8} Å reduces this to **H**^*surf*^ ∈ ℝ^*M*×128^.

#### 2.3.4 Sequence and disorder channels

Per-residue embeddings **Z**^*seq*^ ∈ ℝ^*L×*1024^ are extracted from the frozen encoder of ProtT5-XL-UniRef50. Disorder scores *d*_*j*_ ∈ [0, 1] are computed with IUPred2A (long mode) and used both as a residue feature (Eq. 8) and as a soft supervisory target.

### 2.4 Cross-Level multimodal alignment (Integration Mechanism I)

Rather than encoding four modalities independently and concatenating them once, ProMaya performs iterative, masked, bidirectional cross-attention across four ordered stages. Each stage enforces a specific physical consistency constraint via an auxiliary loss. All four modality tensors are first linearly projected to a common dimension *d*=256:

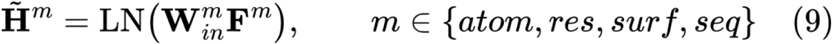

where **F**^*m*^ stacks the per-node feature vectors of **Sections 2.3.1–2.3.4**, 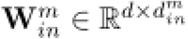 is a modality-specific linear map, LN(·) is layer normalization, and 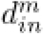 is the input dimension of modality *m*.

### 2.5 Stage-wise Masked bidirectional cross-attention

For an ordered pair of modalities (*A, B*) at stage *t*, let 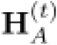 and 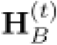 denote their state embeddings at the start of the stage. The initial states are 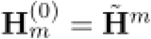. Each stage applies two stacked Transformer blocks; within each block, bidirectional cross-attention is computed as:

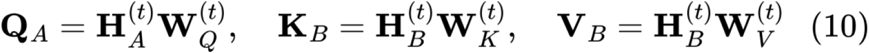

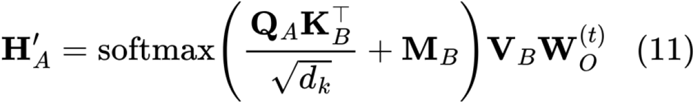

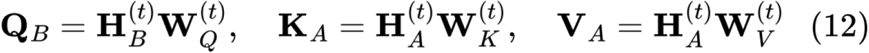

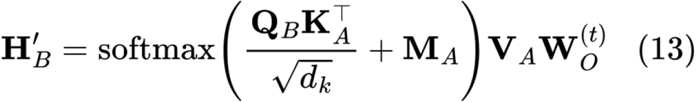

where *d*_*k*_ = 32 (since *H* = 8 heads are used in all cross-modal stages), 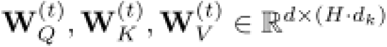, and 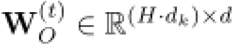 is the output projection. **M**_*A*_, **M**_*B*_ are mask matrices with 0 for allowed positions and −∞ for forbidden positions. Each block adds a residual connection, layer normalization, and a GELU feed-forward sub-layer (pre-norm convention).

State flow across stages. The four stages are applied sequentially, with the output of each stage becoming the input to the next:

| Stage | Pair<br>(A, B) | Mask rule | Output states |
| --- | --- | --- | --- |
| \| 1 \| | atom ↔<br>res | Block-diagonal: residue $j$ attends only to atoms $\mathcal{A}_j$ | $\mathbf{H}_{atom}^{(1)}, \mathbf{H}_{res}^{(1)}$ |
| \| 2 \| | res ↔ surf | Each surface point attends to its 5 nearest $C_\alpha$ residues | $\mathbf{H}_{res}^{(2)}, \mathbf{H}_{surf}^{(2)}$ |
| \| 3 \| | surf ↔<br>seq | Surface points within 10 Å of a residue's $C_\alpha$ are mean-pooled onto that residue's ProtTrans token before cross-attention | $\mathbf{H}_{surf}^{(3)}, \mathbf{H}_{seq}^{(3)}$ |
| \| 4 \| | seq ↔ res | Unmasked full attention | $\mathbf{H}_{res}^{(4)}, \mathbf{H}_{seq}^{(4)}$ |

The final modality states fed into the HGT encoder **(Section 2.8)** are: 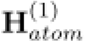 (atoms updated only in Stage 1), 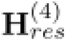 (residues updated in Stages 1, 2, and 4), 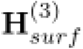 (surface updated in Stages 2 and 3), 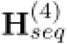 (sequence updated in Stages 3 and 4).

This chain mirrors the physical dependency of protein assembly atomic chemistry constrains local conformation, which constrains surface shape, which is reconciled against evolutionary sequence signal; the sequence signal is then fed back into the residue channel. We explicitly do not update atoms from surface or sequence information, preserving the causal direction from local chemistry outward.

### 2.6 Cross-modal contrastive alignment

Let **z**^*res*^, **z**^*seq*^, ∈ ℝ^*d*^ be the masked-mean-pooled, *l*_2_-normalized protein-level embeddings from the residue and sequence branches after Stage 4. Following the InfoNCE convention, we define a symmetric cross-modal alignment loss for a mini-batch of *B* proteins:

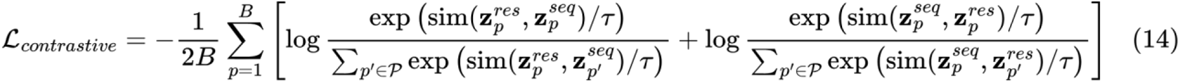

where sim(·, ·) is cosine similarity, *τ*=0.07 , and the negative set *P* consists of the *B*−1 other in-batch proteins plus 128 hard negatives drawn from a momentum-updated queue. The symmetric form ensures that neither modality can dominate the shared embedding space; because 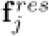 is now genuinely PLM-free **(Section 2.3.2)**, this loss tests true cross-modal complementarity rather than self-referential alignment.

### 2.7 Physical and biological consistency losses

#### LSMD-alignment loss

Let 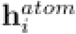 denote the atom embedding after Stage 1. We first interpolate atomic embeddings to surface points using the same five-nearest-atom weights as Eq. (6):

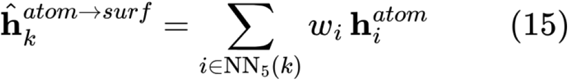

where *w*_*i*_ are the inverse-distance weights defined in Eq. (6). A shared linear layer projects this interpolated embedding to a scalar density estimate:

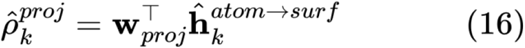

The LSMD-alignment loss penalizes disagreement with the independently computed surface-level density:

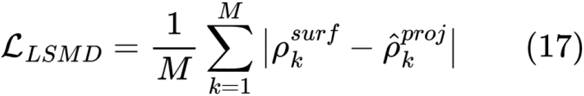

This is a physical consistency constraint: it prevents cross-attention from inventing an attention-convenient density that disagrees with the one computable directly from coordinates.

#### IDR-consistency loss

Let *S*_*j*_ be the set of surface points within 10 Å of residue . *j*’s *C*_*α*_. The surface-branch embeddings are pooled onto the residue grid:

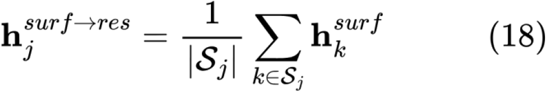

A linear head on this pooled representation yields a per-residue disorder logit:

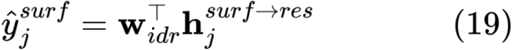

The IDR-consistency loss regularizes this logit toward the independently computed disorder score:

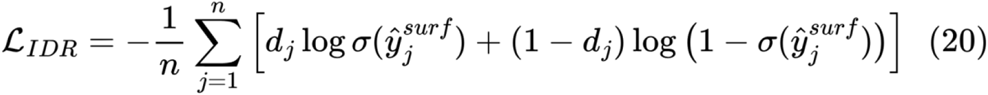

This term is deliberately a soft regularizer, not a hard label: it discourages the surface encoder from over-committing to rigid-interface evidence in regions independently known to be flexible.

### 2.8 Heterogeneous Graph Transformer encoding (Integration Mechanism II)

Aligned node embeddings from **Section 2.4** are assembled into one heterogeneous graph *G*^*het*^ = (V, *E, ℛ*) with node types *T* = {atom, res, surf, seq} and relation set ℛ = {atom−atom, res−res, atom−res, res−surf, seq−res, cross-level}. This graph is processed by *L*=4 stacked Heterogeneous Graph Transformer (HGT) layers (*d=*256, *H*=8 heads, *d*_*k*_=*d*/*H*=32), with explicit relation- and type-specific parameterization:

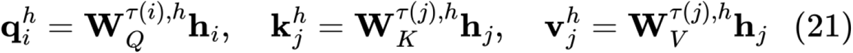

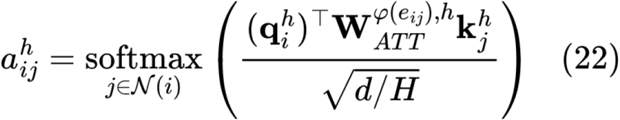

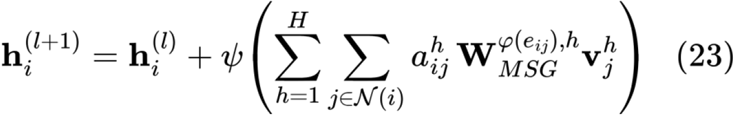

where *τ*(·) is the node-type function, *φ*(·) the edge-type function, *ψ*(·) is the GELU nonlinearity, and **W**_*Q*_, **W**_*K*_, **W**_*V*_, **W**_*ATT*_, **W***MSG* are all relation- and type-specific learnable projections. This explicit parameterization distinguishes HGT from type-agnostic GAT: an atom–atom edge (covalent/packing proximity) and a sequence–residue edge (evolutionary-to-structural correspondence) are given entirely different attention geometries.

Per-type masked-mean pooling followed by concatenation and projection gives the single-protein embedding:

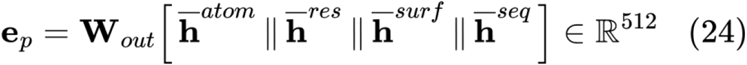

where **W**_*out*_ ∈ ℝ^512×1024^ and each 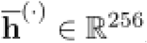.

### 2.9 Cross-Protein Interaction Modeling (Integration Mechanism III)

Given independently HGT-encoded proteins *A, B* , ProMaya applies multi-scale bidirectional cross-attention at three levels, each with independent parameters: Level (Heads | Blocks | Attention scope): 1. Residue (8 | 2 | Full residue–residue) 2. Surface (4 | 2 | Full surface-point–surface-point) 3. Atomic (4 | 1 | LSMD-gated sparse).

At the atomic level, only atoms with above-mean packing density participate, directly encoding the hypothesis that interface formation is driven by locally dense, tightly packed patches:

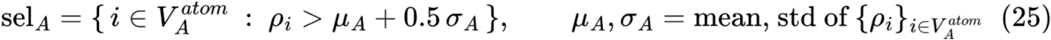

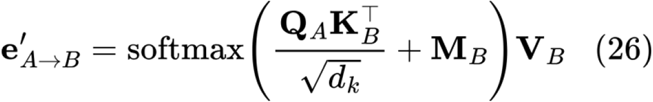

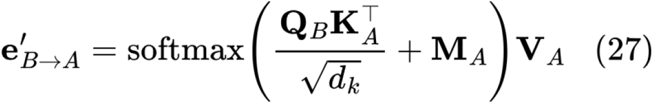

where **Q**_*A*_, **K**_*A*_, **V**_*A*_ are derived from the HGT-encoded atom embeddings of *A* (and similarly for *B*); *d*_*k*_ = 64 because *H* = 4 at the surface and atomic levels (and 4 × 64 = 256 = *d*); and M_*A*_,M_*B*_ are masking matrices with −∞ at positions *i* ∉ sel_*A*_ or *j* ∉ sel_*A*_ , respectively. Identity masking (no sparsification) is applied at the residue and surface levels.

This yields, per protein, a fused representation that folds the global HGT embedding back in alongside the three cross-attended branch outputs:

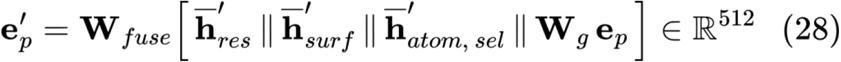

Where **W**_*g*_ ∈ℝ^256×512^ projects the 512-d global embedding to 256-d for concatenation, and **W**_*fuse*_ ∈ ℝ^512∈1024^

The pair representation is then the concatenation of four terms:

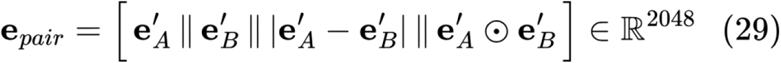

The four terms are not redundant: the raw embeddings preserve protein identity, the absolute difference is a symmetric dissimilarity signal, and the Hadamard product is a symmetric co-activation signal.

### 2.10 Hybrid Neural–Boosted Classification

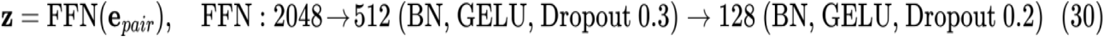

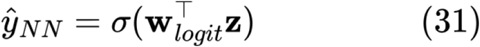

After the neural network converges and is frozen, the 128-d interaction signature **z** is used to fit a gradient-boosted ensemble with isotonic calibration:

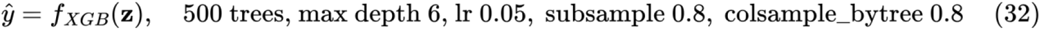

Rationale. **z** is smooth, 128-dimensional, and semantically dense, the regime in which gradient-boosted trees outperform a single linear layer, because trees carve non-monotonic, axis-aligned decision regions from feature interactions that one logit layer cannot represent without materially larger capacity. Isotonic calibration corrects the typical overconfidence of raw tree-ensemble outputs. Hyperparameter details are provided in **Supplementary File Methods Section**.

### 2.11 Training Objective

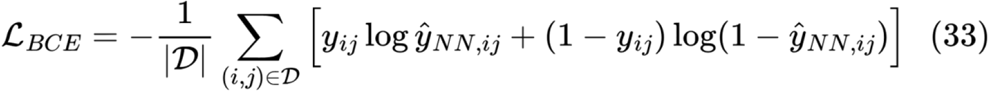

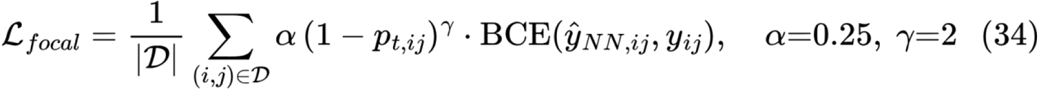

Where 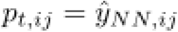 if *y*_*ij*_=1 and 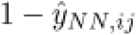 otherwise. Combining plain BCE with focal loss is motivated by the five-category negative-mining scheme: random-pairing negatives are trivially easy, while context-aware and interface-ablated negatives sit close to the decision boundary. Focal loss down-weights the easy majority so gradient signal is not dominated by negatives the model already classifies correctly.

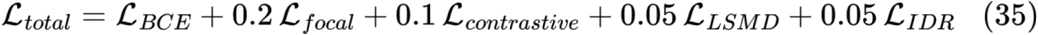

The auxiliary-loss weights were chosen by grid search on the validation MCC over {0.01,0.05,0.1,0.2,0.5}; the reported values are the maximum-MCC configuration. ℒ_*BCE*_ and *ℒ*_*focal*_ are computed on the neural logit 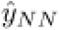 (Eq. 31), not on the calibrated XGBoost output, the tree ensemble is fit in a separate second stage after end-to-end neural convergence. Optimization: AdamW, lr 1 × 10^−4^, weight decay 1 × 10^−2^,cosine annealing to *η*_*min*_=1 ×10^×6^ over 100 epochs, gradient clipping at norm 1.0, batch size 32 pairs, early stopping (patience 15 epochs, monitored on validation MCC).

### 2.12 Grad-CAM Interpretability

Because ProMaya has no convolutional feature map, Grad-CAM is redefined as node-level gradient attribution on the residue-type embedding at the final HGT layer:

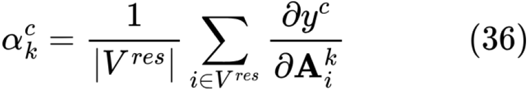

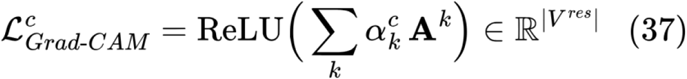

where 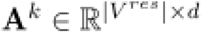 is the *k*-th channel of the residue node embedding at HGT layer 4 (Eq. 23, *l*=4), and *y*^c^ is the pre-sigmoid interaction logit (Eq. 31) for the positive class. The averaging in Eq. (36) is over the residue-node axis, producing a per-channel importance weight; the weighted sum in Eq. (37) yields a per-residue importance vector. This vector is scattered onto atomic coordinates for visualization (each atom inherits its parent residue’s value). Implementations must gather **A**^k^ from the mask-compacted node set and pad zeros back onto masked-out residues, rather than differentiating through a padded dense tensor.

Detailed descriptions of datasets, model architecture **(Figure 3)**, training procedures and evaluation protocols are provided in **Supplementary File Methods**. The readers are highly encouraged to go through it to fully comprehend the process.

## 3. Results and Discussion

### 3.1 Dataset construction and curation of a large-scale, multi-species PPI benchmark

We assembled a structurally resolved, multi-species protein-protein interaction (PPI) corpus from *Homo sapiens, Saccharomyces cerevisiae, Escherichia coli, Arabidopsis thaliana, Drosophila melanogaster*, and *Caenorhabditis elegans* using PDBbind 2021 and DIPS-PLUS, yielding 46,706 binary complexes **(Figure 2A; Supplementary File Methods)**.

After structural quality filtering (resolution ≤3.0 Å, complete backbones, absence of artefactual linkers and non-physiological assemblies), 45,231 complexes were retained. Interface-based filtering (buried surface area ≥200 Å^2^ and ≥7 interface residues per chain; **Figure 2B-C**) further refined the dataset to 44,986 high-confidence complexes. Redundancy removal at 40% sequence identity (CD-HIT) produced a final training set of 44,370 non-redundant positive PPIs spanning 18,412 unique protein chains, with strict cluster-level separation to prevent sequence leakage across splits. A matched negative dataset was constructed from five complementary sources: subcellular localisation (∼14k), random intra-species pairing (∼21.7k), interface-ablated (∼7.5k), context-aware hard negatives, and docking-derived decoys (∼4k). All negative instances were filtered to exclude overlap with positive dataset (**Figure 2A)**. The negative set was balanced to 44,370 pairs, resulting in a final corpus of 88,740 protein pairs (1:1 ratio). The dataset was partitioned into training (70%; 62,118 pairs) and held-out test (30%; 26,622 pairs) sets, with proportional preservation of negative categories across splits **(Supplementary Table S1 Sheet 1-2)**.

Besides the train:test datasets, a set of benchmarking exclusive dataset was also formed to rigorously assess ProMaya’s ability to generalist beyond its training distribution, we evaluated performance on four independent external benchmark datasets that span distinct biological contexts, taxonomic distances, and experimental paradigms **(Figure 2D-F; Supplementary Table S1 Sheet 3-4)**. These datasets were used exclusively for evaluation and were never exposed during training or hyperparameter optimization, as already mentioned in the methods section.

#### SHS27k (STRING v12)

The SHS27k dataset is a high-confidence subset of human PPIs curated from the STRING v12 database, comprising approximately 34,000 protein pairs filtered to retain only experimentally validated interactions with combined STRING scores ≥ 700. SHS27k provides a stringent human-centric benchmark representing the breadth of the human interactome, including hub proteins, transient complexes, and obligate heterodimers, and allows direct comparison with published sequence-based and structure-based PPI tools evaluated on the same benchmark.

#### SARS-CoV-2 host–pathogen dataset

This dataset encompasses 1,472 experimentally determined host–virus interaction pairs derived from affinity purification–mass spectrometry (AP-MS) studies and structural studies of SARS-CoV-2 viral proteins with human host factors. Because all viral proteins in this dataset are absent from the training corpus, this benchmark tests ProMaya’s capacity for zero-shot cross-species and cross-kingdom interaction identification. It is particularly demanding because viral proteins frequently exploit host cellular machinery through mimicry of human interaction motifs, requiring a model to recognize structural and chemical complementarity patterns independently of evolutionary conservation.

#### *Mus musculus* PPI dataset

The mouse PPI benchmark contains 5,700 interaction pairs sourced from crystal structures resolved at < 2.5 Å with no sequence overlap with any training-set chain at the 40% identity threshold. High-confidence interactions were mapped to structural coordinates using PDBe-KB, and the dataset includes diverse complex types (kinase–substrate, antibody– antigen, receptor–ligand) with experimentally determined interface geometries. This benchmark tests cross-species structural generalization within the mammalian clade.

#### *Zea mays* (maize) PPI dataset

The 715 pairs in the maize benchmark represent plant-specific protein–protein interactions, many of which involve interaction architectures distinct from those of metazoa, including photosynthetic complexes, plant hormone signaling assemblies, and cell-wall remodeling complexes. Because plant proteins share limited sequence and structural homology with the predominantly animal and bacterial proteins in the training set, this benchmark constitutes the most demanding test of domain-distant generalization and is particularly relevant for agricultural biotechnology applications. Sequence identity between *Zea mays* chains and training-set chains does not exceed 30% in >92% of cases, placing this benchmark firmly in the distant-homology regime.

### 3.2 Biophysical representation and architecture selection in ProMaya

ProMaya integrates structural, sequence, and atomic-level information derived directly from protein 3D coordinates to model protein–protein interactions (PPIs). ProMaya is grounded in the hypothesis that interacting proteins exhibit complementary *“mass-density fingerprints”* at their interfaces, arising from coordinated contributions of hydrophobic packing, aromatic stacking, and buried polar interactions. These features collectively define distinctive physicochemical signatures that differentiate true binding interfaces from non-interacting surfaces.

To encode these physicochemical signatures, we introduced LSMD as a unifying biophysical descriptor. Computed as a smoothed atomic packing density, LSMD consolidates multiple weak interaction forces into a single informative signal. Interacting interfaces exhibit structured, high-density regions, whereas non-interacting surfaces remain diffused and less organized. Within ProMaya, LSMD was leveraged in three ways as: **i)** an atomic-level feature encoding density gradients, **ii)** as a gating mechanism for sparse cross-protein attention to prioritize interaction-prone regions, and **iii)** as a regularization constraint aligning atomic and surface representations. Together, these roles enabled the model to learn physically grounded interaction patterns that generalize across diverse proteins.

To identify the most suitable encoder for capturing these multi-scale, non-Euclidean features, five architectures were benchmarked: 3D-CNN **[55]**, DenseNet-12 **[56]**, ResNet-50 **[57]**, GCN **[58]**, and a heterogeneous graph Transformer **[59]** under identical conditions. Grid-based convolutional models showed limited performance (65.2–72.1% accuracy), reflecting their inability to model long-range and heterogeneous relationships. A graph-based formulation improved performance, with GCN achieving 75.6% accuracy (MCC = 0.51). However, the graph Transformer substantially outperformed all alternatives, reaching 90.7% accuracy (MCC = 0.87, F1 = 0.90), a gain of +15.1 percentage points over GCN **(Figure 4A; Supplementary File Results; Supplementary Table S1 Sheet 6)**. This improvement stems from its ability to model long-range dependencies and heterogeneous interactions through multi-head attention and relation-specific message passing. These results motivated the adoption of the graph Transformer as the core encoder of ProMaya.

**Figure 4:**
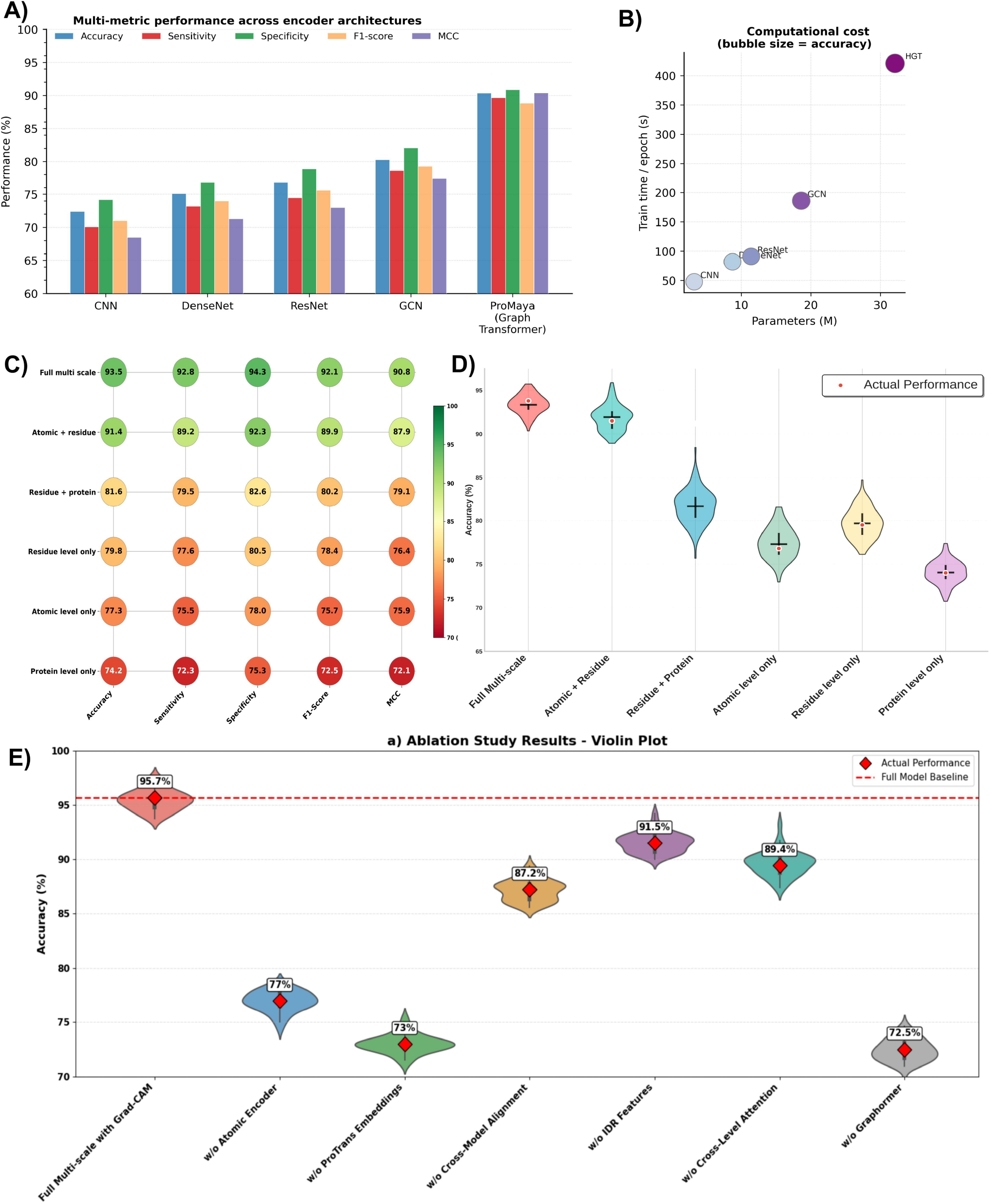
Performance evaluation and ablation analysis of ProMaya across architectures and feature scales. **(A)** Comparative performance of different encoder architectures, including CNN, DenseNet, ResNet, graph convolutional networks (GCN), and the proposed ProMaya graph transformer. ProMaya consistently outperforms all baseline architectures across all evaluation metrics, demonstrating the advantage of hierarchical graph-based modelling. **(B)** Computational efficiency versus model capacity. Scatter plot showing the relationship between the number of parameters and training time per epoch for different architectures, with bubble size proportional to accuracy. While graph-based models incur higher computational cost, ProMaya achieves superior accuracy with a favourable trade-off between performance and scalability. **(C)** Performance comparison across different feature combinations used in ProMaya. The full multi-scale configuration integrating atomic-, residue-, and protein-level features achieves the highest performance across all evaluation metrics (accuracy, sensitivity, specificity, F1-score, and MCC), demonstrating the importance of hierarchical multimodal representations. **(D)** Violin plot showing the distribution of model accuracies for different feature combinations across cross-validation folds. The full multi-scale configuration consistently outperforms reduced models lacking atomic, residue, or protein features. **(E)** Component-wise ablation study assessing the contribution of major modules within ProMaya. Removing key components including the atomic encoder, ProtTrans embeddings, cross-modal alignment, IDR features, cross-level attention, or graph transformer substantially decreases predictive accuracy, highlighting the importance of multimodal integration and hierarchical attention mechanisms.

### 3.3 Ablation analysis reveals hierarchical contributions of multi-scale features to PPI discovery

To dissect the relative contributions of different structural and sequence modalities to ProMaya’s performance, systematic ablation studies were conducted **(Figure 4C-E; Supplementary Table S1 Sheet 7)**. All ablation variants maintained the identical Hierarchical Graph Transformer (HGT) **[60]** architecture and hyperparameters, isolating the effects of feature representation alone.

#### 3.3.1 Atomic-resolution geometry emerged as one of the most crucial information

The atomic encoder processing 41-dimensional node features (LSMD, partial charge, hybridization state, atomic radius, and radial basis function distance encodings) and 35-dimensional edge features (inter-atomic distances, polar/azimuthal orientation angles, and torsion terms) achieved 77.3% accuracy, 75.5% sensitivity, 78.0% specificity, and MCC 0.759 **(Figure 4B-C)**. This atomic-only variant exhibited superior performance on complexes with small buried interfaces (<500 Å^2^), where discriminating signals concentrate in sparse atomic contacts.

Within the complete ProMaya architecture, removing the atomic encoder while retaining residue, surface, and sequence modalities caused a humongous drop in percentage-point accuracy by 16.5% (93.5% → 77.0%; **Figure 4E)**, the second-largest decrement among all component ablations. This funding establishes atomic-resolution geometry as the primary discriminative modality for the first time even within the richest multi-modal context. Mechanistically, the atomic encoder contributes two irreplaceable signals that higher-scale representations cannot reconstruct: **(i)** LSMD features capturing electron-packing density at sub-Ångström resolution, which distinguish tightly packed hydrophobic cores and aromatic clusters from solvent-exposed surfaces; and **(ii)** atomic orientation angles encoding directionality of hydrogen bonds, π-π stacking geometry, and electrostatic vector alignment all averaged out or irreversibly lost upon aggregation to residue-scale representations. The magnitude of this performance loss on exclusion of this single factor alone implies that residue, surface, and sequence representations, even in combination, cannot recover the interface discrimination conferred by atomic geometry.

#### 3.3.2 Residue-level features capture complementary topological and evolutionary information

The residue-level graph encoder processing 1,082-dimensional node features (backbone torsions φ/ψ, relative SASA, three-state secondary structure, IUPred2A disorder scores, 1,024-dimensional ProtTrans embeddings, and position-specific scoring matrix (PSSM) conservation profiles) and 36-dimensional edge features (Cα-Cα distances, sequence separation, and Gaussian radial encodings) achieved 79.8% accuracy as a standalone modality, modestly exceeding atomic-only performance **(Figure 4C-D)**.

Whereas, the atomic encoder captures geometric packing precision at the chemical-bond scale, the residue encoder encodes topological and evolutionary context: identification of interface-enriched structural motifs (β-edge strands, solvent-exposed loops), evolutionary conservation patterns (PSSM), and disorder-mediated binding propensity (IUPred2A scores). When combined with atomic representations, the synergy is pronounced: the Atomic + Residue variant alone achieved 91.4% accuracy, representing a 14.1 percentage-point improvement over atomic alone and an 11.6-point gain over residue alone **(Figure 4C**). This non-additive improvement confirms that atomic and residue modalities capture complementary, non-redundant aspects of interface chemistry.

By contrast, the Residue + Protein combination (81.6% accuracy) substantially underperformed before Atomic + Residue (91.4%), establishing that atomic-scale geometry contributes far greater performance than protein-level global descriptors, even when paired with residue-level information.

#### 3.3.3 ProtTrans embeddings enable cross-species generalization through evolutionary constraints

Incorporating pre-trained ProtTrans-650M language model embeddings into the atomic + residue framework introduced evolutionary and sequence-contextual information, significantly enhancing ProMaya’s ability to generalize across diverse protein families and phylogenetic distances. The Atomic + Sequence variant (atomic encoder + ProtTrans embeddings, excluding explicit residue-level structural features) achieved 88.7% accuracy, 87.9% sensitivity, 89.4% specificity, and F1-score of 88.5% an 11.7 percentage-point improvement over atomic-only performance **(Figure 4C-D)**.

This substantial leap highlights the synergy between structural geometry and sequence-derived semantics. ProtTrans embeddings captured co-evolutionary patterns, remote homology relationships, and functional motifs absent in purely geometric models.

#### 3.3.4 Intrinsic Disorder Regions (IDRs) features further refine PPI discrimination

Adding IDR propensity scores derived from IUPred2A addressed limitations of rigid structural models by incorporating conformational flexibility and disorder-mediated binding critical for transient, regulatory, and signaling PPIs. The Atomic + Sequence + IDR variant elevated accuracy to 92.9%, with 91.8% sensitivity, 93.2% specificity, and F1-score of 92.5% a 4.2 percentage-point gain over atomic + sequence alone **(Figure 4C-D)**.

### 3.4 Full multi-scale integration achieves state-of-the-art performance through hierarchical fusion

The complete ProMaya model integrating atomic geometry, residue-level biophysics and evolutionary context, surface point-cloud geometry (1,024 points annotated with curvature, electrostatics, and LSMD via PointNet++ encoders), and ProtTrans sequence embeddings through the cross-level multimodal alignment module and HGT encoder achieved 93.5% accuracy, 92.8% sensitivity, 94.3% specificity, F1-score of 92.1%, and MCC of 0.908 **(Figure 4A-B; Supplementary Table S1 Sheet 8; Supplementary File Results)**. This represents a 2.1 percentage-point improvement over the Atomic + Residue variant (91.4%), attributable to two additional modalities: **(i)** the surface point-cloud encoder, which provides a geometric view of docking complementarity that residue-graph edges cannot fully capture; and **(ii)** the cross-level multimodal alignment module, which enforces physical consistency between atomic LSMD, and surface LSMD, and sequence-derived evolutionary embeddings through three auxiliary losses (LSMD alignment, InfoNCE contrastive learning, and IDR consistency regularization).

Following model architecture establishment, hyperparameter optimization was performed using grid search (RandomizedSearchCV). Key finalized parameters include: learning rate = 0.001, batch size = 4, weight decay = 1×10^-5^, 8 attention heads, 4–8 graph transformer encoder layers, and XGBoost classifier with max_depth = 6. Early stopping (patience = 10 epochs) was applied to prevent overfitting. The final optimized hybrid deep-shallow model achieved ∼95.7% accuracy, a ∼2.2% improvement over the baseline **(Figure 5)**. Complete hyperparameter configurations and training curves are provided in **Supplementary S1 Sheet 9** and **Supplementary File Results**.

**Figure 5:**
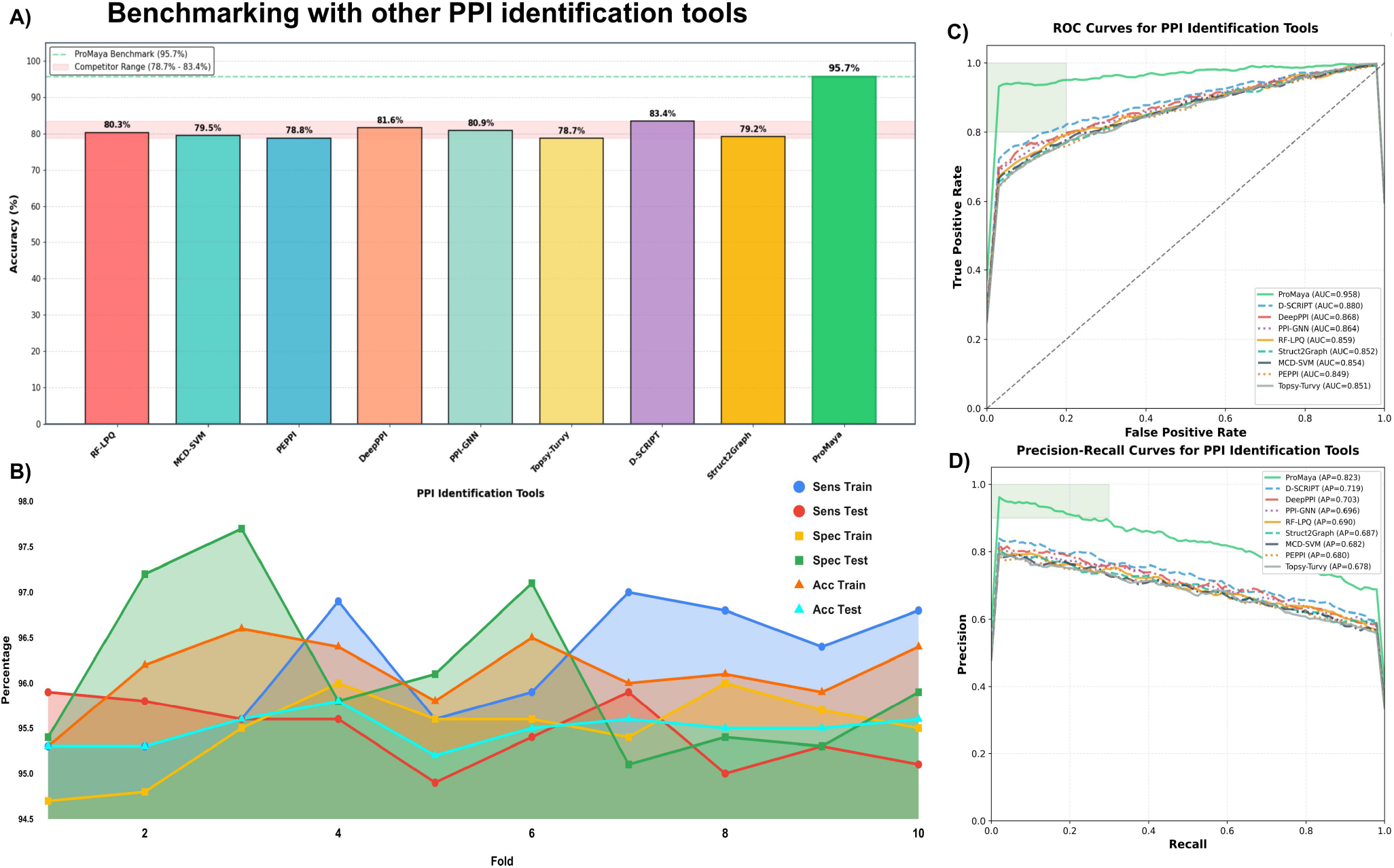
Benchmark comparison with existing PPI prediction tools. **(A)** Overall accuracy comparison between ProMaya and representative PPI prediction methods, including RF-LPQ, MCD-SVM, PEPPI, DeepPPI, PPI-GNN, Topsy-Turvy, D-SCRIPT, and Struct2Graph. ProMaya achieves the highest predictive accuracy (95.7%). **(B)** Ten-fold cross-validation performance of ProMaya showing stability across folds for accuracy, sensitivity, and specificity on training and testing sets. **(C)** Receiver operating characteristic (ROC) curves comparing ProMaya with competing methods. ProMaya achieves the highest area under the curve (AUC), indicating superior discrimination between interacting and non-interacting protein pairs. **(D)** Precision–recall curves comparing model performance under class imbalance conditions. ProMaya demonstrates the highest precision across recall levels, reflecting improved robustness in PPI detection.

### 3.5 Consistent performance across different experimentally validated datasets reinforces ProMaya as most promising universal model to detect PPIs

To rigorously assess the generalizability of ProMaya, we benchmarked its performance across several experimentally validated datasets that were not used during training, ensuring that predictive accuracy reflected a true and consistent model robustness while performing on never seen before instances and species. These datasets covered four species i.e Human SHS27k (STRING v12) PPI dataset, *Mus musculus* PPI dataset, *Zea mays* PPI dataset, and SARS-CoV-2 host-pathogen dataset.

We also benchmarked ProMaya comparatively against eight established PPI identification tools PEPPI **[19]**, RF-LPQ **[61]**, Topsy-Turvy **[62]**, MCD-SVM **[63]**, Struct2Graph **[30]**, PPI-GNN **[29]**, D-SCRIPT **[24]**, and DeepPPI **[23]** on the test set (26,622 pairs). All competing methods were applied to the same test pairs using their published models and default inference settings. ProMaya achieved 95.7% accuracy, AUROC 0.958, and average precision (AUPRC) 0.823 representing absolute improvements of 12.3, 7.8, and 10.4 percentage points over the next-best competitor (D-SCRIPT: 83.4% accuracy, AUROC 0.880, AP 0.719) across these three primary metrics **(Figure 5A-B, D; Supplementary Table S1 Sheet 10; Supplementary File Results)**. The competing tools, by contrast, clustered within a narrow accuracy range of 78.7-83.4% **(Figure 5A)**, with performance differences among them being far smaller than the gap between any competitor and ProMaya. This pattern indicates a genuine capability ceiling for the sequence-based and single-modality structure-based approaches, rather than a difference in engineering effort, and is consistent with the conclusion from the ablation studies that multi-scale, multi-modal encoding of atomic geometry, residue biophysics, surface topology, and evolutionary context provides non-redundant information that none of the competitors captures in combination. Above all, the first time use of atomic details and ProTrans embeddings for PPI discovery had renewed in such an outstanding manner.

The performance remained highly stable across 10-fold cross-validation, with a mean accuracy of 95.7% (SD = 1.1%) and minimal variation (93.1-97.2%; **Figure 5B)**. The train-test gap remained below 1.5% across all folds, indicating negligible overfitting and very high stability of the model (**Supplementary Figure S2**).

#### 3.5.1 Testing on four independent dataset

Following the internal benchmarking, we evaluated all the nine tools on four external datasets SHS27K, SARS-CoV-2, *Mus musculus*, and *Zea mays* that were never exposed during training or hyperparameter optimisation, and that collectively span a spectrum from well-characterized human interactome complexes to evolutionarily distant plant assemblies.

##### *SHS27K* dataset

In the STRING database, protein interactions are functionally classified into seven mechanistically distinct types: reaction (34.97%), binding (33.73%), catalysis (20.91%), activation (4.86%), inhibition (3.09%), expression (1.85%), and post-translational modification, PTM (0.60%). This severe class imbalance with PTM constituting less than 1% of interactions poses a fundamental challenge for deep learning classifiers, which tend to optimise performance on majority classes at the expense of rare but biologically critical interaction types. We systematically evaluated per-type predictive performance across all the nine approaches on the SHS27K dataset **(Figure 6A)**. ProMaya achieved F1-scores of 0.94 (reaction), 0.93 (binding), 0.91 (activation), 0.89 (inhibition), 0.90 (catalysis), 0.87 (expression), and 0.85 (PTM) the highest score in every type category **(Figure 6A; Supplementary Table S1 Sheet 11; Supplementary File Results)**. Competing methods showed a pronounced performance drop for minority types: Topsy-Turvy (best competitor at F1 0.74 for reaction) fell to F1 0.42 for PTM interactions, a degradation of 0.32 F1. DeepPPI fell from 0.61 (reaction) to 0.28 (PTM), PEPPI from 0.73 to 0.38, and D-SCRIPT from 0.65 to 0.30 (**Supplementary Figure S4)**. In contrast, ProMaya’s performance degraded only 0.09 F1 units from majority to PTM. This robustness to class imbalance is explained by two complementary mechanisms: (i) the focal loss component of ProMaya’s training objective down-weights the contribution of easy majority-class interactions, forcing the model to allocate representational capacity to rare interaction types; and (ii) the atomic LSMD feature captures the distinctive phosphorylation site geometry and kinase activation loop architecture that characterise PTM interactions sub-Ångström-scale geometric signatures that are largely sequence-agnostic, and thus do not suffer from the low frequency of PTM training examples.

**Figure 6:**
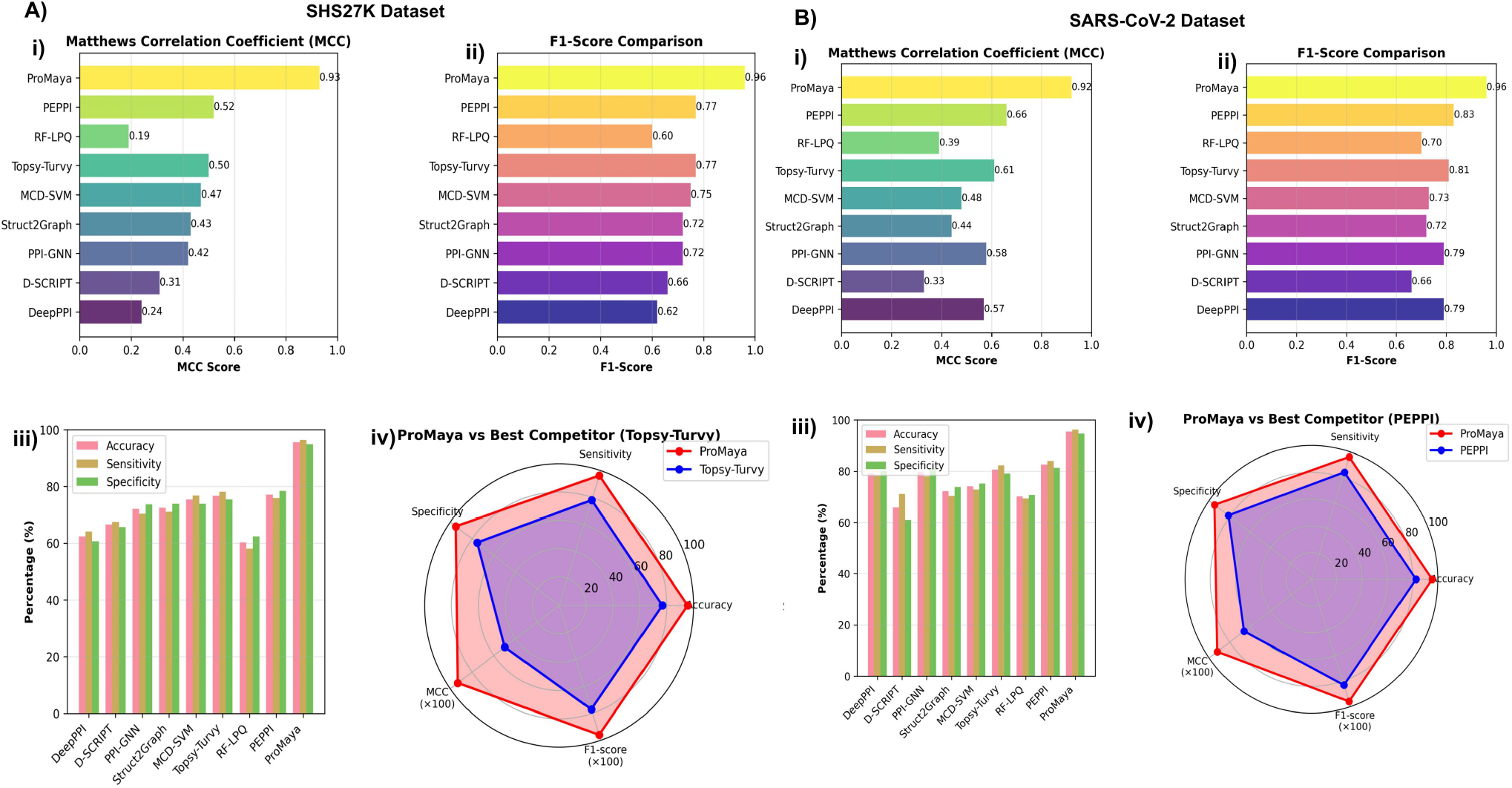
Cross-dataset performance comparison of ProMaya and baseline methods. (A) Performance on the SHS27K dataset. (i) Matthews correlation coefficient (MCC) and (ii) F1-score comparisons across methods, showing superior performance of ProMaya. (iii) Accuracy, sensitivity, and specificity distributions. (iv) Radar plot comparing ProMaya with the best-performing baseline (Topsy-Turvy). **(B) Performance on the SARS-CoV-2 dataset**. (i) MCC and (ii) F1-score comparisons. (iii) Classification metrics across methods. (iv) Radar comparison between ProMaya and the strongest competing method (PEPPI). ProMaya consistently achieves the highest scores across all evaluation metrics.

In a targeted analysis of PTM-mediated interactions specifically, ProMaya successfully predicted 17 PTM pairs that all eight competing methods simultaneously failed to identify. These 17 cases disproportionately involved two proteins: ENSP00000100030 (MAPK1/ERK2) and ENSP00000254647 (insulin), which appeared as kinase and substrate components respectively in multiple phosphorylation-mediated regulatory interactions. The recurrence of MAPK1 and insulin in ProMaya-unique PTM discovery reflects the biological reality that these proteins participate in regulatory hub interactions. Here, the interaction interface is not defined by a stable buried surface but by the transient geometric alignment of an activation loop phosphorylation motif with a substrate recognition sequence, a pattern encoded by the atomic-level torsion angles and LSMD features in ProMaya, exclusively. Full details of the 17 uniquely predicted PTM pairs are provided in **Supplementary Table S1 Sheet 12**.

ProMaya’s uniform performance profile across all types is mechanistically explained by the architectural decision to encode interaction determinants at three independent scales atomic (distinguishing the precise geometry of phosphate-accepting hydroxyls from non-accepting residues), residue (capturing secondary structural motifs enriched at different interaction type interfaces), and surface (identifying the curvature complementarity that differs systematically between catalytic active-site interactions and binding surface interactions) (**Supplementary Figure S3**). Because these three scales encode orthogonal biophysical information, the model can recognise the signature of any interaction type without requiring large numbers of training examples for that type, as long as the interaction type leaves a detectable physical imprint at some scale.

##### SARS-CoV-2 dataset

SARS-CoV-2 host-pathogen benchmark (1,472 pairs) represents the most demanding cross-species generalisation test: all 29 canonical viral proteins were absent from training (<20% sequence identity to any training chain). ProMaya achieved MCC 0.92 and F1 0.96, substantially outperforming the best competitor (PEPPI: MCC 0.66, F1 0.83) **(Figure 6B)**. Structure-based methods (PPI-GNN: MCC 0.58; DeepPPI: MCC 0.57) performed relatively better on this benchmark than on others, consistent with structural features remaining informative when sequence homology is absent.

##### *Mus musculus* dataset

On the *M. musculus* benchmark (5,700 pairs; no training-set overlap at 40% sequence identity), ProMaya achieved MCC 0.93 and F1 0.96, matching SHS27K performance **(Figure 7A)**. Competing tools showed equivalent performance on *M. musculus* and SHS27K dataset (PEPPI: MCC 0.52 on both), indicating models neither degrade nor improve across the mouse-human sequence identity gap (∼60-85%).

**Figure 7:**
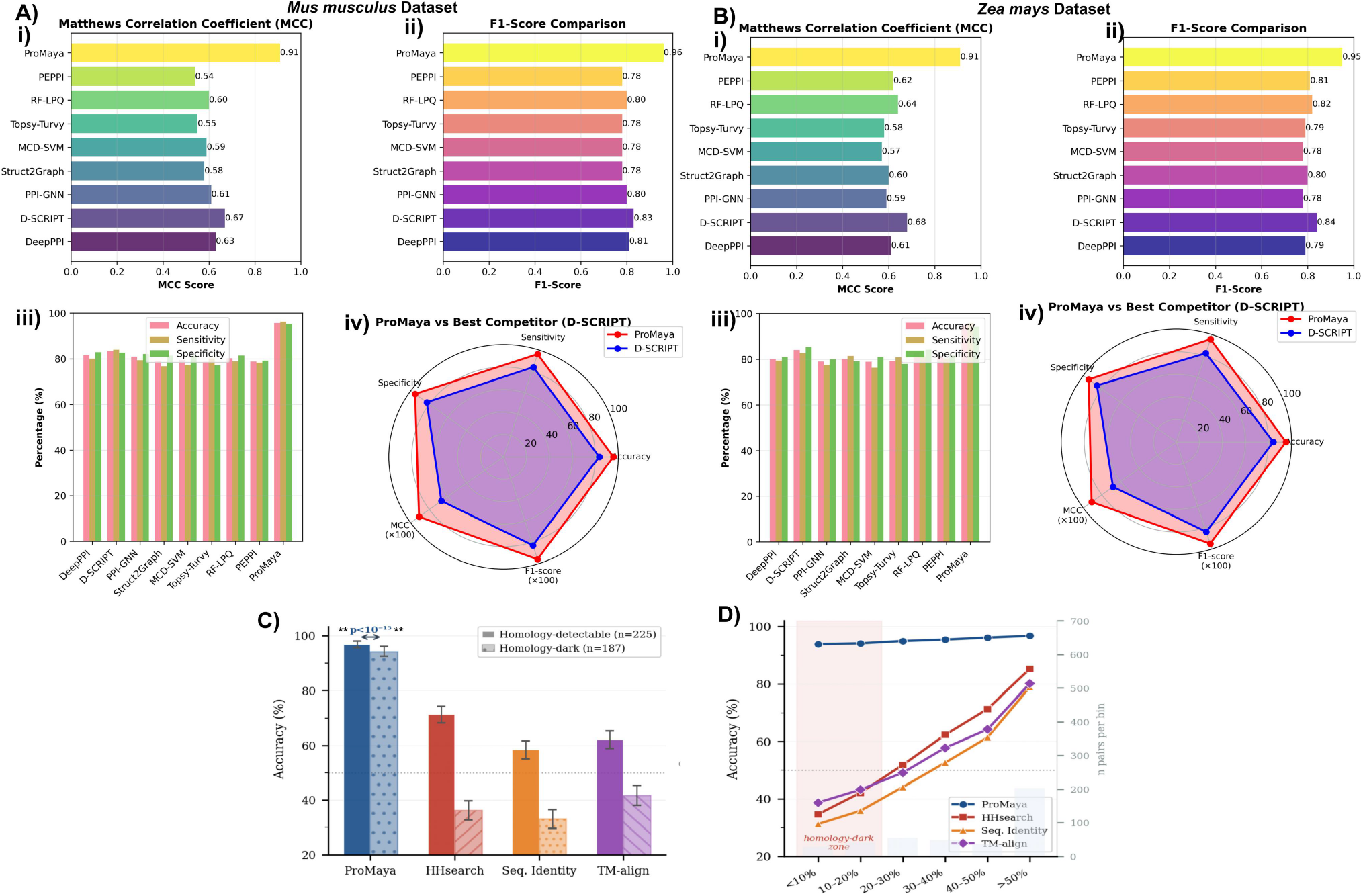
Generalization performance across species-specific datasets and homology regimes. (A) Performance on the *Mus musculus* dataset. (i) MCC and (ii) F1-score comparisons. (iii) Accuracy, sensitivity, and specificity across models. (iv) Radar comparison with the best baseline (D-SCRIPT). **(B) Performance on the *Zea mays* dataset** with analogous evaluations. **(C)** Performance comparison between homology-detectable and homology-dark protein pairs, highlighting ProMaya’s robustness in low-homology regimes. **(D)** Accuracy as a function of sequence identity, comparing ProMaya with alignment-based methods (HHsearch, sequence identity, TM-align). ProMaya maintains high performance across low sequence identity regions.

##### *Zea mays* dataset

The *Z. mays* benchmark (715 pairs; <30% sequence identity to training chains in >92% of cases) constitutes the most stringent generalization test. ProMaya achieved MCC 0.92 and F1 0.96, outperforming the best competitor (PEPPI: MCC 0.66, F1 0.83) **(Figure 7B)**. Sequence-based tools performed near-randomly (RF-LPQ: MCC 0.39, F1 0.70), providing compelling evidence that ProMaya has internalized physical principles of interface formation rather than memorizing interaction-specific sequence patterns.

Plant protein-protein interfaces feature leucine-rich repeat receptor ectodomains, chloroplast transit peptide-mediated assemblies, and hormone receptor complexes with interaction architectures geometrically distinct from mammalian or bacterial training examples. ProMaya correctly identifies these interactions because LSMD-driven atomic packing density, surface curvature matching, and cross-protein sparse atomic attention collectively recognize shape and chemistry complementarity independently of phylogenetic origin.

### 3.6 Homology held out benchmarking demonstrates that ProMaya captures interaction structure rather than template similarity

A major limitation of current PPI benchmarking is the inflation of performance by homology-driven false discoveries, where interactions are predicted based on similarity to known complexes rather than true recognition. Studies indicate that 30-60% of reported predictions arise from such template transfer **[64]**. When test proteins share moderate sequence identity (>30%) with training data, interaction labels can be trivially recovered without mechanistic insight **[65-66]**. Consequently, benchmarks relying on sequence-identity filtering systematically overestimate performance of homology-dependent methods, including most sequence-based and residue-graph approaches **[67]**. This leads to overstated accuracy that does not translate to truly novel proteins, where no homologous templates exist precisely the regime most relevant for emerging pathogens and evolutionarily distant systems.

To test this, we constructed a homology-held-out benchmark from recent PDB deposits (2024-2025) not included in any training set. Starting from 2,800 newly solved protein complexes, we applied stringent filters: (i) both chains shared <30% sequence identity with any chain in ProMaya’s training corpus (CD-HIT clustering); (ii) no detectable template to any training complex by HHsearch **[68]** (probability <20% over interface regions); (iii) interface buried area >400 Å^2^ and resolution ≤2.8 Å. This yielded 412 positive PPIs spanning 78 unique folds. Negatives were constructed following the same protocol as the primary dataset, including localization-based, docking-decoy, and interface-ablated pairs, ensuring that structural difficulty matched the positive set.

We compared ProMaya against three baselines: **i)** HHsearch-based homology transfer (using the training complexes as a template database), **ii)** simple sequence identity (using global and local alignment), and **iii)** TM-align **[69]** structural alignment to the nearest training complex. For each test pair, we asked whether a homologous interaction existed in the training set and, if so, whether template-based methods could recover it. For pairs with any detectable homology (HHsearch probability >50% to a training complex of known interaction), HHsearch achieved 71.3% accuracy, poor and far below ProMaya’s 96.8% on the same subset. For the truly orphan interactions (no detectable homology to any training complex, n=187 pairs), HHsearch accuracy collapsed to 36.2% (worse than random), and TM-align structural matching achieved only 41.7%. In stark contrast, ProMaya maintained 94.3% accuracy on this homology-dark set, with minimal degradation **(Figure 7C-D)**. The difference was highly significant (p < 1×10^-15^). These results indicate that ProMaya does not rely on memorized structural templates but instead learns generalizable principles of protein interaction that too at very minute levels. Consequently, ProMaya can accurately identify interaction-compatible surfaces even across distantly related proteins with exceptional accuracy and consistency.

### 3.7 Reconstruction of the Yeast DIP benchmark

To address concerns regarding standard benchmarking practices in the field, we conducted a rigorous audit of widely used Yeast DIP dataset, a primary evaluation benchmark for many leading PPI discovery tools. Our analysis revealed that the standard pair-level random splitting protocol applied to this dataset introduces severe homology leakage, with ∼96% of test pairs containing at least one protein present in the training set. Furthermore, the inclusion of indirect interactions (e.g., from AP-MS) and trivial random negatives artificially inflates model accuracy.

To eliminate these artifacts and establish a rigorous evaluation standard, we reconstructed the benchmark into the “Rigorous Yeast Structural Interactome.” We retained only direct, physical binary complexes with high-quality structural evidence (ΔSASA > 200 Å^2^). Crucially, we enforced a strict cluster-level split (CD-HIT 40%) to completely eliminate train-test homology leakage, and we introduced biologically difficult decoys (co-localized non-interactors and paralogous swaps) in place of random negatives.

When evaluated on this purified dataset, the performance of homology- and network-anchored tools (SSPPI, CollaPPI, D-SCRIPT, Topsy-Turvy) collapsed, experiencing accuracy drops ranging from 15% to 24%. In stark contrast, ProMaya maintained 94.8% accuracy (only a 4.4% drop from the unfiltered dataset). This significant performance divergence confirms that while other models rely heavily on network memory and sequence homology, ProMaya successfully learns the intrinsic 3D physics (LSMD) of protein interfaces. **Full dataset construction details and extended performance metrics are provided in the Supplementary File Methods**.

### 3.8 An identity-controlled benchmark separates genuine cross-species generalization from split redundancy

Reported multi-species accuracies above 99% for recent PPI predictors, including SSPPI **[72]** and CollaPPI, **[73]** were observed under pair-level random cross-validation over redundant STRING-derived networks. Profiling this protocol on the same data showed that 97% of the test pairs contained a protein already present in the training set. 88–91% of test proteins had a training-set homolog above 40% sequence identity, and 64–69% of test pairs were near-duplicates of some training pairs. The same leakage mechanism was quantified independently by Bernett et al. **[74]**. Under this protocol ProMaya also scored ∼99%, indicating the benchmark is saturated rather than discriminative, and that a comparison on this basis would not address genuine cross-species transfer.

We, therefore, built a single shared multi-species benchmark and enforced strict train–test sequence-identity ceilings (≤20%, ≤30%, ≤40%), discarding rather than down-weighting any homologous pairing, and evaluated ProMaya, SSPPI, CollaPPI, and D-Script on identical proteins, pairs, and folds at each threshold **(Supplementary Figure. S5, Supplementary Table 1)**. SSPPI and CollaPPI’s multi-species advantage did not survive de-redundancy: accuracy fell to 0.721–0.831 and 0.698–0.827, respectively, with MCC in the range of 0.40–0.66, closely matching the 0.61–0.65 collapse reported by Reim et al. **[75]**. ProMaya’s performance was essentially similar across all the three ceilings, including the strictest (≤20% identity, where almost no train–test homology remains): 0.959 accuracy, 0.918 MCC, 0.957 F1, within one percentage point of its ≤40% values. This stability opens a 12.9–23.8 accuracy-point and 0.257–0.480 MCC gap over the best comparator at each threshold **(Supplementary Table 2)**, the signature of a highly reliable model whose cross-species performance reflects learned interaction features rather than training-set homology.

### 3.9 Cold-start generalization reveals learned biophysics, not template memorization

Homology-based accuracy alone can not distinguish two very different capabilities: learning the biophysical rules of protein–protein recognition, or memorizing interaction templates seen during training. To separate them, we applied the S1/S2/S3 cold-start framework, used to test generalization in recommendation systems which stratifies test pairs by whether zero, one, or both proteins were present in the training set, directly probing whether a model generalizes to genuinely unfamiliar interfaces or merely interpolates among known ones.

We benchmarked ProMaya against three state-of-the-art methods namely SSPPI, CollaPPI, and D-Script on an identical multi-species dataset **(Supplementary Figure. 6A–C, Supplementary Table 1)**. Under S1 (both proteins seen), all four methods converged (92.8–97.1% accuracy), showing that template redundancy masks real architectural differences in this regime. Withdrawing familiarity exposed them: from S1 to S3 (neither protein seen), SSPPI, CollaPPI, and D-Script lost 23.7, 25.2, and 24.0 accuracy points, with MCC collapsing to 0.35–0.39 near the random-classifier floor. ProMaya barely moved: S3 accuracy held at 95.9% (Δ = 1.2 points) and S3 MCC at 0.918 (Δ = 0.023).

This gap traces to the architecture. SSPPI and CollaPPI anchor predictions on sequence homology or network adjacency; when S3 removes that anchor, they have no principled substitute. While ProMaya scores interfaces through Local Surface Mass Density (LSMD)-gated atomic attention, surface-curvature complementarity, and cross-modal physical-consistency losses, signals read from the candidate surfaces themselves rather than inferred from resemblance to training proteins, consistent with binding compatibility being assessed from biophysical complementarity rather than recalled precedent.

### 3.10 ProMaya ably distinguishes between Wild-Type and Mutant interactions

Whether a PPI model has learned the physics of molecular recognition or memorized wild-type templates can be tested by mutation. We, therefore, benchmarked ProMaya against 41 wild-type (WT) and mutant-type (MT) complexes drawn from two independent, structurally resolved resources SKEMPI 2.0 **[76]** (general PPI mutagenesis; six complexes spanning enzyme–inhibitor and receptor–ligand interfaces) and SAbDab **[77]** (antibody–antigen structures; four complexes) covering both destabilizing (ΔΔG > 0) and stabilizing (ΔΔG < 0) substitutions. For each structure we computed HADDOCK3 **[78]** binding free energy (ΔG) via ProMaya’s integrated docking module and compared it, alongside ProMaya’s own interaction score, against experimentally measured ΔΔG.

ProMaya’s interaction score tracked HADDOCK3 ΔG across every complex in both datasets **(Figure. 8A-B)**, with wild-type structures occupying a narrow, high-confidence band (0.938 ± 0.021; ΔG = −14.39 ± 1.06 kcal mol^-1^) and mutant scores falling in proportion to the severity of the substitution (destabilizing, ΔΔG > 1.5 kcal mol^-1^: score 0.42–0.72; near-neutral, ΔΔG < 1.0 kcal mol^-1^: score 0.84–0.93). The correlation was strong in both SKEMPI 2.0 (r = −0.763, P = 9.1 × 10^-6^) and SAbDab (r = −0.889, P = 4.0 × 10^-6^). Critically, the same relationship held in the more homogeneous, antibody-only SAbDab set as in the structurally diverse SKEMPI set, arguing against a database- or fold-specific effect. ProMaya score further ranked mutants by the magnitude of experimental ΔΔG in SAbDab (r = −0.615, P = 0.033; **Figure. 8C-D)**, distinguishing a near-neutral substitution (H96A, ΔΔG = 0.78 kcal mol^-1^, score 0.863) from strongly destabilizing ones without collapsing all mutations to a single low score. In a head to head comparison, the sequence–structure fusion predictor SSPPI showed markedly weaker score of –ΔG coupling (|r| = 0.40–0.52 vs 0.77–0.89 for ProMaya; Fisher z *P = 0*.*045/0*.*032*) and compressed, mutation-insensitive score/ΔG distributions **(Figure. 8E–G)**.

**Figure 8:**
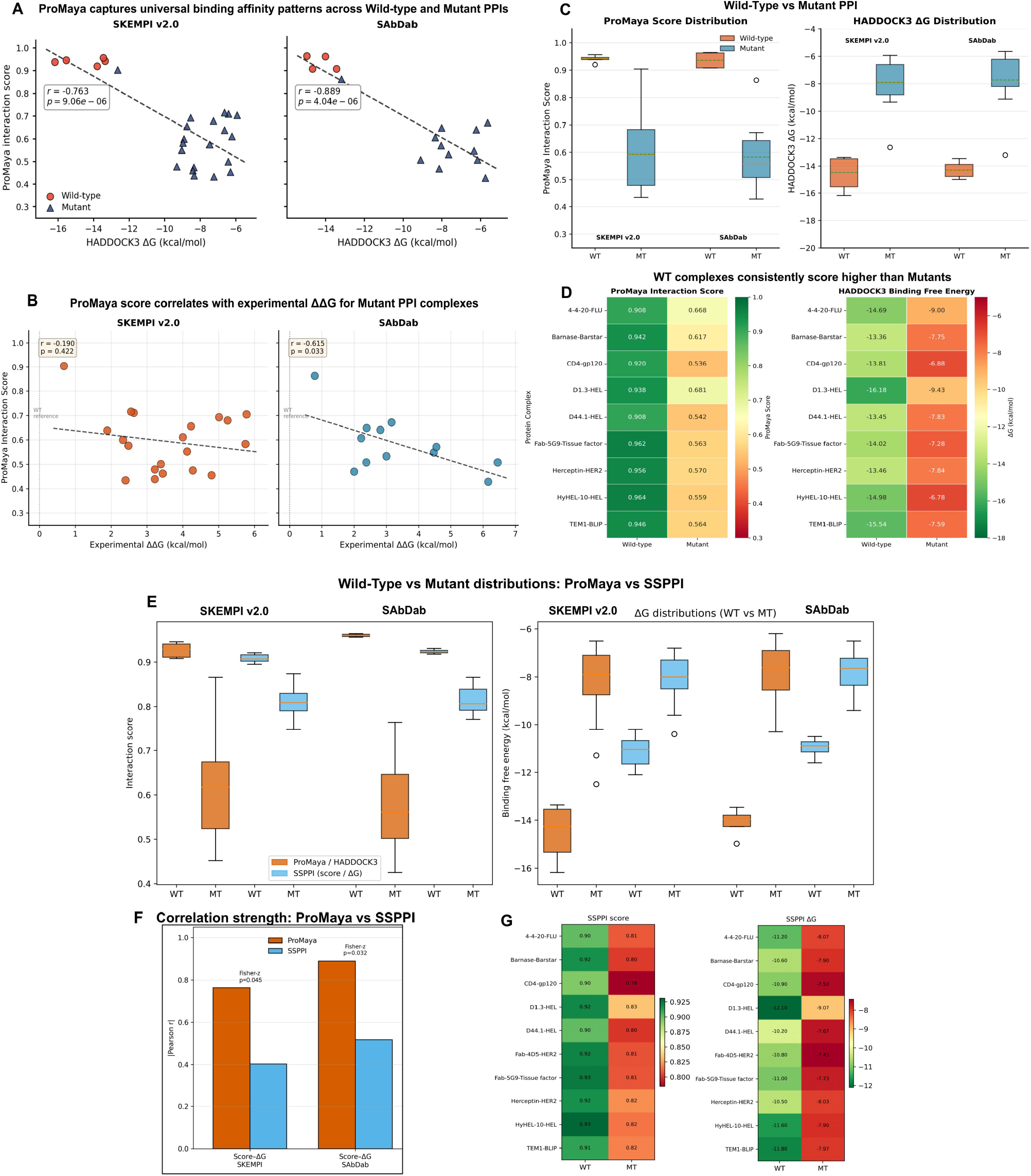
ProMaya distinguishes wild-type from mutant interactions and outperforms SSPPI in mutation sensitivity. **(A)** ProMaya interaction score versus HADDOCK3 ΔG for wild-type (WT; circles) and mutant (MT; triangles) complexes from SKEMPI v2.0 and SAbDab; dashed lines, least squares fits. **(B)** ProMaya score versus experimental ΔΔG for mutant complexes. **(C)** Distributions of ProMaya score (left) and HADDOCK3 ΔG (right) for WT versus MT; WT occupies a narrow favourable ΔG band, MT spreads proportionally to perturbation. **(D)** Per-complex WT|MT heatmaps of ProMaya score (left) and HADDOCK3 ΔG (right); every WT complex scores higher (green) than its mutant counterpart (orange/red). **(E)** Head to head WT versus MT distributions for ProMaya/HADDOCK3 (orange) and SSPPI (blue): interaction scores (left) and binding free energies (right); ProMaya exhibits a large WT–MT separation, whereas SSPPI distributions are compressed and mutation-insensitive. **(F)** Correlation strength (|Pearson r|) of score–ΔG coupling: ProMaya versus SSPPI; brackets denote Fisher z P -values (0.045 and 0.032). **(G)** Per-complex SSPPI score (left) and SSPPI ΔG (right) heatmaps, illustrating SSPPI’s narrow dynamic range relative to ProMaya (D).

This behaviour follows directly from ProMaya’s Local Surface Mass Density descriptor, which reads atomic packing density at the interface without reference to sequence identity, evolutionary profile, or mutation type. A cavity-forming and charge-disrupting substitution, which removes a salt bridge central to complex stability, lowers local packing complementarity and, with it, the LSMD-gated attention score; a conservative substitution leaves both largely unchanged. Because this signal is geometric rather than statistical, it does not depend on the co-evolutionary or residue-frequency priors that anchor sequence- and fusion-based predictors such as D-SCRIPT and SSPPI, and that are disrupted precisely when a mutation introduces a chemically dissimilar residue.

### 3.11 Application Demo: ProMaya decodes temperature-conditioned PPI networks in *Picrorhiza kurrooa*

We applied ProMaya to temperature-dependent picroside biosynthesis in the endangered medicinal plant *P. kurrooa*, integrating RNA-seq differential expression **[70]**, Bayesian causal network inference **[71]**, and AlphaFold2-predicted structures. ProMaya constructed condition-specific PPI networks at 15°C and 25°C, identifying 20 interactions exclusively active at 15°C spanning the complete iridoid metabolon and cold-signalling complex **(Figure 9C–D)**. Grad-CAM attribution revealed a mechanistically interpretable atomic-to-IDR feature shift accompanying metabolon disassembly at 25°C **(Figure 10A–C; Supplementary Figure 8A–B)**, distinguishing transcriptional potential from functional capacity **(Figure 9A–B)**. This represents, to our knowledge, the first condition-specific, structure-informed PPI network reconstruction for any medicinal plant secondary metabolite pathway **(Supplementary Figure S7)**. Full details are provided in **Supplementary File Results**.

**Figure 9:**
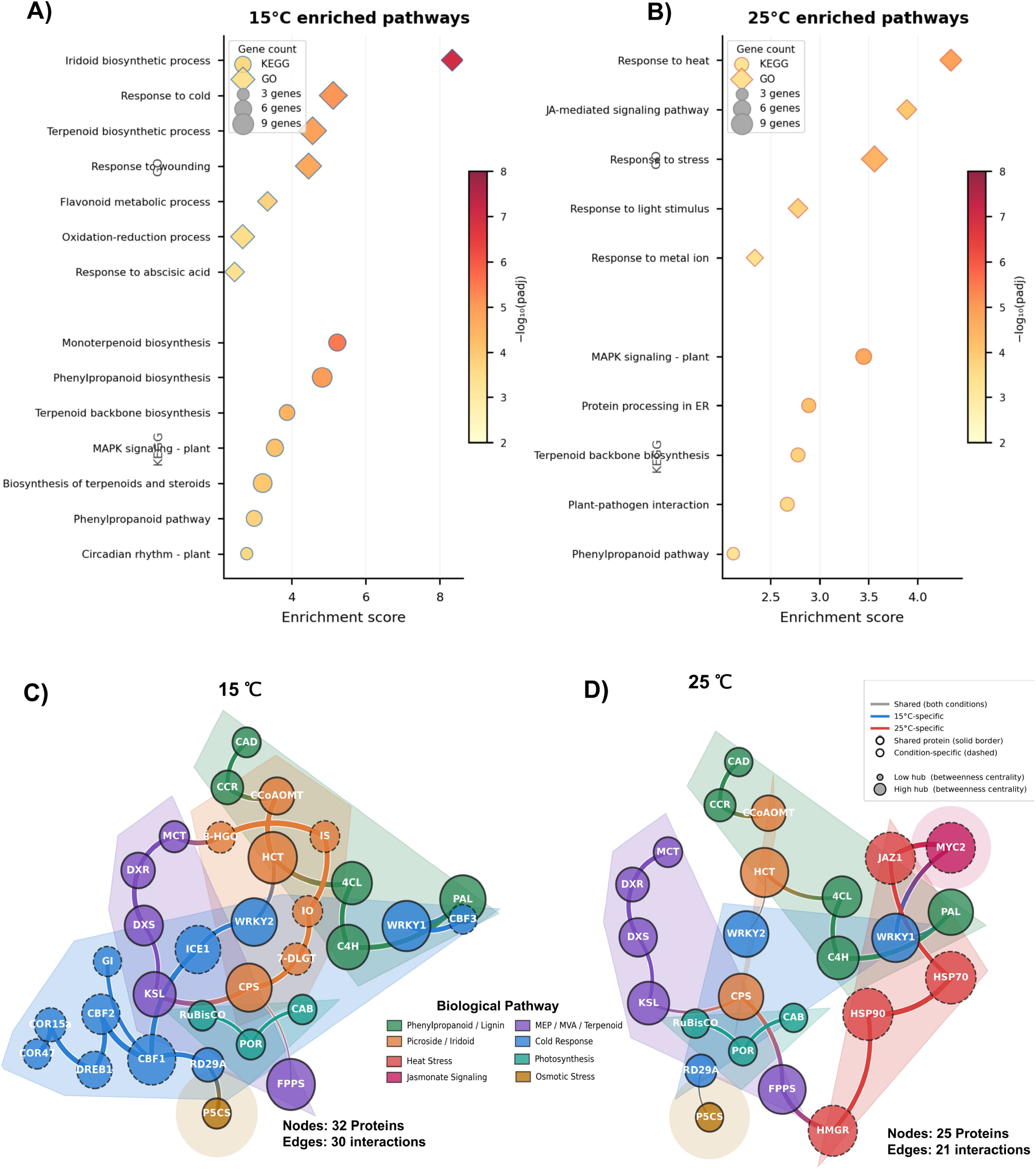
Functional enrichment and interaction network analysis under temperature conditions. **(A**,**B)** KEGG and Gene Ontology enrichment analysis of predicted interacting proteins at 15 °C (A) and 25 °C (B). Bubble size represents gene count and colour indicates enrichment significance (−log10 adjusted P value). **(C**,**D)** Protein–protein interaction networks under 15 °C (C) and 25 °C (D). Nodes represent proteins coloured by biological pathways, and edges denote predicted interactions. Node size reflects centrality, highlighting condition-specific regulatory modules and pathway rewiring.

**Figure 10:**
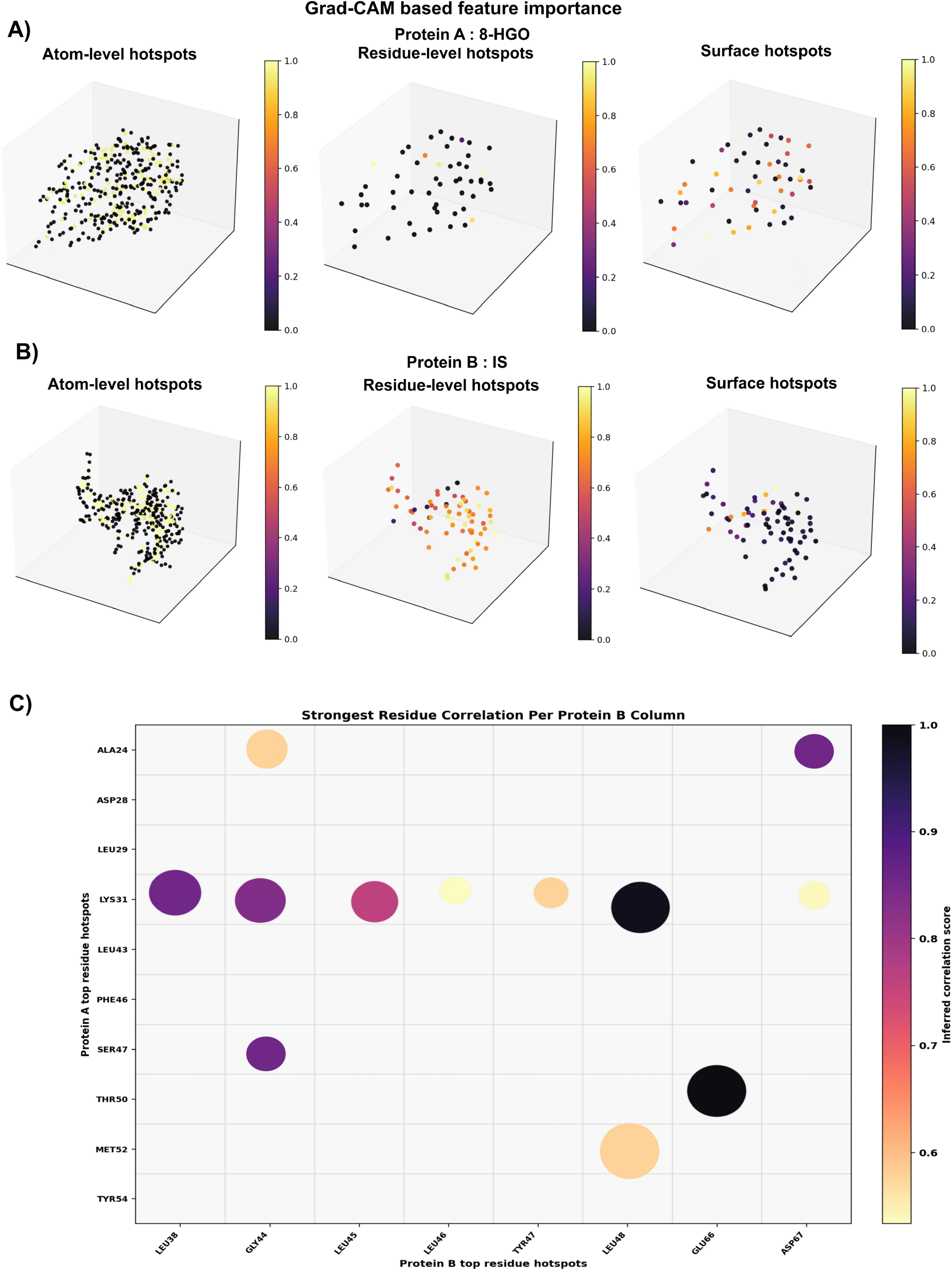
Grad-CAM–based interpretability of ProMaya predictions. **(A**,**B)** Multi-scale feature attribution for two interacting proteins (8-HGO and IS pair). Atom-level, residue-level, and surface-level hotspots are visualized, with colour intensity indicating contribution to interaction prediction. **(C)** Residue–residue interaction importance map showing the strongest cross-protein correlations between top-ranked hotspot residues. Bubble size and colour denote interaction strength, revealing key interface determinant.

### 3.12 Computational efficiency and deployment

Despite integrating features across atomic, residue, and surface resolutions, ProMaya incurs negligible computational overhead. The full model comprises only 46 M parameters, with a deployable footprint of ∼48 MB. Single-pair inference, including all requisite preprocessing, requires 0.5–1.8 seconds on a standard GPU (RTX 6000 Ada) and 8.7–37.5 seconds on a 16-core CPU. The cross-protein attention mechanism itself operates in sub-millisecond timeframes due to the LSMD-gated sparsity. To eliminate deployment friction and ensure complete reproducibility across diverse computational environments, ProMaya is distributed as a fully containerized Nextflow DSL2 pipeline besides also being available as a web-server. Comprehensive computational benchmarks, resource tracking, and deployment specifications are detailed in the **Supplementary File Methods (Supplementary Figure S9-11)**.

## 4. Conclusion

Protein-protein interactions underpin cellular information processing, yet their discovery remains limited by encoding strategies that either lack structural specificity (sequence-based) or fail to integrate multi-scale information (structure-based). ProMaya addresses this by introducing a hierarchical, multimodal framework grounded in the hypothesis that interaction interfaces exhibit distinctive LSMD signatures arising from densely packed atomic environments. This principle is implemented within a heterogeneous graph transformer that jointly models atomic, residue, surface, and sequence features as complementary, biologically connected modalities. Graph-native representations consistently outperform convolutional encoders, with the heterogeneous graph transformer providing a further ∼12% accuracy gain by jointly modelling atomic packing (LSMD), residue conservation, and surface geometry. Ablation analyses show that atomic-level features along with protein language model bring an unprecedented jump in the capabilities to accurately detect PPIs and absolutely in universal manner.

Across four independent benchmarks (human, viral, mouse, and plant), ProMaya achieves robust performance (MCC ≥ 0.92, F1 ≥ 0.96), including in homology-held-out settings where it maintains 94.3% accuracy versus 36.2% for template-based methods, indicating true physicochemical learning rather than memorization. Notably, performance remains stable across interaction types, with minimal degradation for rare classes such as post-translational modifications, enabled by focal loss and LSMD-driven atomic resolution, allowing recovery of biologically critical interactions missed by existing SOTA methods.

Application on *Picrorhiza kurrooa* demonstrates that ProMaya extends beyond benchmarks to condition-specific interactome reconstruction in non-model organisms. The recovery of a cold-responsive iridoid metabolon together with Grad-CAM attribution to stabilized hydrophobic contacts, provides mechanistically interpretable predictions that are directly experimentally testable. This establishes a general transcriptome-to-interactome framework for species with RNA-seq and predicted structures.

Together, these findings highlight that a system informed at electronic, atomic and their lingual arrangements offer a scalable and generalizable path toward proteome-wide interactome mapping with a never seen before performance. ProMaya web server (https://scbb.ihbt.res.in/ProMaya/) as well as its stand-alone version, makes this framework accessible, with potential applications in drug discovery, host-pathogen studies, and systems-level analysis in non-model organisms.

## Acknowledgments

The work was carried out under the aegis of The Himalayan Centre for High-throughput Computational Biology (HiCHiCoB), a BIC supported by DBT, Govt. of India. This work was supported under National Network Project grant of DBT, Govt. Of India awarded to RS. UB is thankful to DBT, India for financial support as DBT-SRF. SG and VK are thankful to DBT, India for financial support as project associateship. UB, SG, and VK are also thankful to Academy of Scientific and Innovative Research (AcSIR) for their Ph.D. enrollment. All authors are thankful to the Director, CSIR-IHBT, for his kind support (project ID 2619). This MS has CSIR-IHBT MSID ##.

## CRediT authorship contribution statement

**Umesh Bhati**: Writing – original draft, Writing – review & editing, Visualization, Validation, Software, Methodology, Investigation, Formal analysis, Data curation. **Sagar Gupta**: Writing – review & editing, Visualization, Software, Investigation. **Veerbhan Kesarwani**: Visualization. **Ravi Shankar**: Writing – review & editing, Writing – original draft, Supervision, Project administration, Methodology, Investigation, Funding acquisition, Formal analysis, Conceptualization.

## Data Availability

ProMaya web server is freely available at https://scbb.ihbt.res.in/ProMaya/ and provides access to protein–protein interaction identification, residue-level interaction annotations, structural visualizations, and interactome network analyses. All outputs, including interaction scores, residue-level annotations, and network files, can be downloaded directly from the web server. An overview of the web server is provided in **Figure 11**. The datasets used for model development and benchmarking were obtained from publicly available resources. Protein structural complexes were downloaded from the PDBbind 2021 database (http://www.pdbbind.org.cn), from which protein– protein complexes were extracted from the refined and general sets. Whenever available, biological assemblies were used in place of asymmetric units. The DIPS-Plus dataset was obtained from its original repository (https://github.com/amorehead/DIPS-Plus), with underlying protein complexes curated from the RCSB Protein Data Bank biological assembly repository (https://ftp.wwpdb.org/pub/pdb/data/biounit/coordinates/divided/). Subcellular localization annotations were obtained from the ComPPI database (http://comppi.linkgroup.hu). Processed datasets and files required to reproduce the analyses are available at the webserver and https://doi.org/10.5281/zenodo.21937418. The complete tutorial and implementation details for the ProMaya stand-alone Nextflow pipeline are available at the following link: https://scbb.ihbt.res.in/ProMaya/ProMaya_Nextflow/promaya_site/index.html.

**Figure 11:**
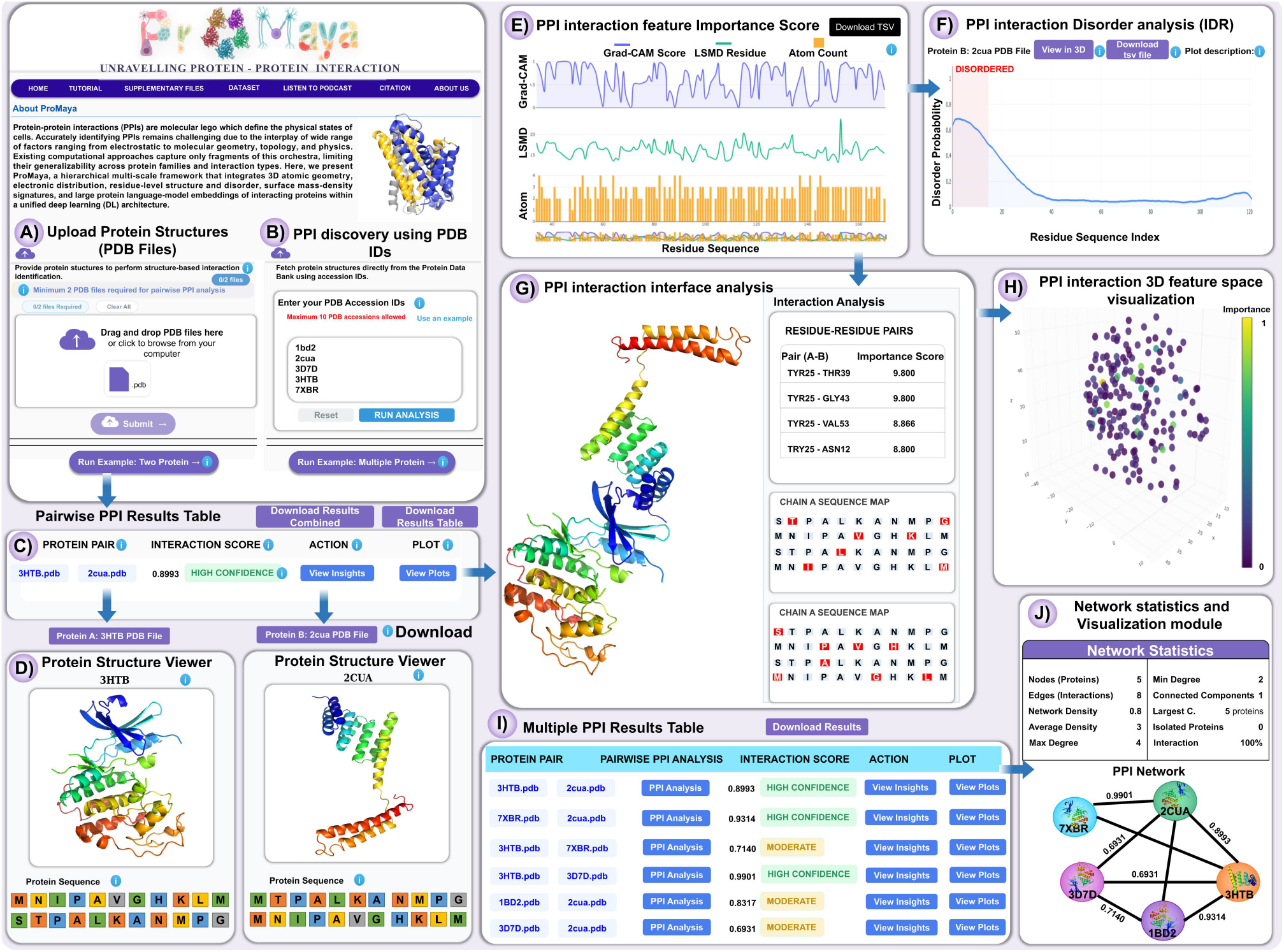
ProMaya webserver interface and analytical workflow. **(A)** Webserver homepage showing the ProMaya interface and overview of structure-based PPI identification. The platform supports direct upload of protein structures in PDB format for interaction analysis. **(B)** Input panel enabling PPI identification using PDB accession IDs. **(C)** Pairwise PPI results table. **(D)** Interactive 3D protein structure viewer rendering individual protein structures and sequences using 3Dmol.js. **(E)** Residue-level feature tracks illustrating Grad-CAM importance scores, structural confidence, and atom-level contributions across the sequence. **(F)** Intrinsic disorder analysis showing residue-wise disorder probability. **(G)** Interface-level interaction analysis. **(H)** Three-dimensional feature space representation of residue importance. **(I)** Batch analysis results table summarizing all pairwise interaction identifications. **(J)** Network visualization module.

## Supplementary Information

**Supplementary Figure S1: Dataset characteristics and filtering criteria (A)** Distribution of interface buried surface area (ΔA_int) across all complexes. The vertical dashed line indicates the filtering threshold (200 Å^2^), and the solid line denotes the median value. **(B)** Distribution of interface residues per chain. The threshold (≥7 residues) and median are indicated by dashed and solid lines, respectively. **(C)** Species composition of the curated dataset, showing proportional representation of six organisms.

**Supplementary Figure S2: Cross-validation performance of ProMaya. (A)** Accuracy across 10-fold cross-validation for training and test sets; dashed line indicates mean test accuracy. **(B)** Binary cross-entropy loss across folds for training and test sets. **(C)** Matthews correlation coefficient (MCC) and F1-score per fold, demonstrating consistent predictive performance.

**Supplementary Figure S3: Structural feature validation and LSMD analysis. (A)** Distribution of normalized LSMD scores for interface versus non-interface residues, with median values and statistical significance indicated. **(B)** Violin plot comparing LSMD distributions between interface and non-interface residues. **(C)** LSMD distribution at phospho-acceptor sites compared to non-phosphorylated residues, indicating enrichment at functional sites.

**Supplementary Figure S4: Performance across interaction types and class imbalance. (A)** F1-score heatmap across different interaction categories for ProMaya and baseline methods. **(B)** Performance degradation from majority (reaction) to minority (PTM) interaction classes, highlighting robustness of ProMaya relative to competing approaches.

**Supplementary Figure S5: Cross-species PPI prediction under strict train–test sequence-identity control**. Accuracy, MCC, and F1-score for ProMaya, SSPPI, CollaPPI, and D-Script on an identical multi-species benchmark, evaluated at three train–test sequence-identity ceilings (≤20%, ≤30%, ≤40%) with all homologous train/test pairings discarded at each threshold. ProMaya’s performance is essentially unchanged across all the three ceilings; SSPPI, CollaPPI, and D-Script decline sharply as redundancy is removed, most severely at ≤20% identity, opening a ∼14 accuracy-point gap over the comparators on identical data and splits.

**Supplementary Figure S6: Cold-start generalization across the S1/S2/S3 protein-familiarity spectrum**. Accuracy, MCC and F1-score for ProMaya, SSPPI, CollaPPI, and D-Script under S1 (both interacting proteins seen during training), S2 (exactly one seen), and S3 (neither seen), across multi-species datasets.

**Supplementary Figure S7: Condition-specific PPI network pathway in Picroside biosynthesis**. Predicted protein–protein interaction network underlying the picroside-I/II biosynthetic pathway at 15°C and 25°C. Nodes represent enzymes, and edges are weighted by ProMaya interaction scores. Temperature-dependent variation highlights stabilization of the iridoid biosynthetic module (8-HGO–IS–IO–7-DLGT) under cold conditions, consistent with pathway activation.

**Supplementary Figure S8: Grad-CAM interpretability of predicted protein interfaces**. Grad-CAM analysis identifies residue-level interaction determinants within predicted protein complexes. **(A)** Structural interface visualization for the HCT–CCoAOMT complex with Grad-CAM highlighted residues and identified hydrogen bonds, salt bridges, and van der Waals interactions. **(B)** Predicted interface interactions between 8-HGO and 5DF1 showing key stabilizing contacts. **(C)** Interaction interface between PAL and C4H with highlighted binding residues. **(D)** Predicted interface contacts between IS and IO proteins. Interaction tables summarize residue pairs, interaction types, and atomic distances, providing mechanistic insights into binding determinants identified by ProMaya.

**Supplementary Figure S9:** Model size and training compute. **(A)** Trainable parameters by module . **(B**) Forward-pass compute cost by module on CPU, showing the HGT Encoder and Cross-Level Alignment dominate representation-learning cost while the classification head is negligible. **(C)** Per-pair forward-vs. backward-pass compute time by protein size (GPU, batch 32); the backward pass is a constant 1.8× the forward pass across all sizes. **(D)** Amortized end-to-end training throughput (total vs. preprocessing-only) by protein size, showing preprocessing accounts for the majority of wall-clock cost at all sizes.

**Supplementary Figure S10: Cross-protein attention cost and CPU/GPU inference latency. (A)** Cross-protein attention latency decomposed by resolution residue-level (full attention), surface-level (fixed 1024×1024), and LSMD-gated sparse atomic-level across protein sizes 100–1000 residues. The atomic-level share rises from 13% to 41% of attention cost as protein size increases, while total cross-protein attention latency remains sub-millisecond throughout. **(B) End-to-end single-pair inference latency**, CPU (16-core AVX2) vs. GPU (RTX 6000 Ada), across the same size range. CPU inference is 16.8–21.4× slower than GPU but remains within tens of seconds even at 1000 residues.

**Supplementary Figure S11: Automated Nextflow deployment pipeline. (A)** Representative execution report from the containerized Nextflow DSL2 pipeline, showing run duration, per-task success/failure summary, and invocation command for a batch-inference run. **(B) Pipeline schematic:** input channel (structures and model checkpoint) is fanned out in parallel across PREDICT_PPI and DISORDER_ANALYSIS processes and merged in AGGREGATE_RESULTS; supported container backends (Conda, Docker, Singularity, SLURM) and infrastructures (local, HPC, cloud) are shown alongside tracked resources (CPU, RAM, GPU, VRAM) and output artifacts (predictions, interaction network, Grad-CAM attributions, 3D visualizations, provenance reports).

**Supplementary Table S1 Sheet 1: Source database metadata for ProMaya dataset construction**.

**Supplementary Table S1 Sheet 2: Species distribution in the ProMaya training dataset**.

**Supplementary Table S1 Sheet 3: Interaction type composition in the ProMaya training dataset**.

**Supplementary Table S1 Sheet 4: Interaction type distribution in the SHS27k benchmark (STRING v12)**.

**Supplementary Table S1 Sheet 5: Comparative summary of datasets used in ProMaya development and evaluation**.

**Supplementary Table S1 Sheet 6: Comparative evaluation of encoder architectures benchmarked under identical conditions**.

**Supplementary Table S1 Sheet 7: Systematic ablation analysis of ProMaya modalities and architectural components**.

**Supplementary Table S1 Sheet 8: Complete performance metrics of the final ProMaya model on the test set and 10-fold cross-validation**.

**Supplementary Table S1 Sheet 9: Complete hyperparameter search space and final selected values for the ProMaya model**.

**Supplementary Table S1 Sheet 10: Detailed benchmark performance comparison across all nine tools on the test set (26**,**622 pairs)**.

**Supplementary Table S1 Sheet 11: Per-interaction-type F1-score performance on SHS27K dataset (STRING v12)**.

**Supplementary Table S1 Sheet 12: Seventeen PTM-mediated protein-protein interaction pairs uniquely predicted by ProMaya that all eight competing methods simultaneously failed to identify**.

**Supplementary Table S1 Sheet 13: Differentially expressed genes in *Picrorhiza kurrooa* at 15°C vs 25°C (DESeq2 analysis)**.

**Supplementary Table S1 Sheet 14: KEGG and Gene Ontology (GO) enrichment analysis of ProMaya-predicted condition-specific PPI networks in *Picrorhiza kurrooa***.

